# The neural geometry of naturalistic surprise: Graded boundary breakthroughs in the human brain

**DOI:** 10.64898/2026.09.03.748562

**Authors:** Yunqiang Hao, Jingyi Li, Chao Jiang

**Affiliations:** School of Psychology, Capital Normal University, Beijing 100048, China

**Keywords:** surprise, neural state transition, whole-brain geometry, boundary breakthrough, naturalistic cognition

## Abstract

How whole-brain neural states reorganize when events violate observers’ predictions remains unclear. Using naturalistic magic-viewing fMRI, we show that more surprising events carry the whole-brain neural state progressively farther beyond the range occupied during non-surprise viewing. This graded departure is accompanied by coordinated reconfiguration across sensory and association systems rather than uniform response amplification. We call this phenomenon a boundary break-through. To quantify its depth, we used non-surprise brain states to define a reference range along the surprise-related transition direction. Boundary exceedance (*E*_BB_) expresses the depth beyond this range in bits; each additional bit means that a reference state is half as likely to reach at least that far. On this scale, depth increased with independently rated surprise intensity, specifically in the breakthrough state and not in the paired reference state. Peri-event trajectories showed that the event deepened an excursion already under way. Exploratory analyses linked exceedance to one-week recognition memory in a surprise-dependent manner. Together, these findings identify a graded systems-level relationship between surprise intensity and the departure of the whole-brain neural state from its non-surprise reference range.

## Introduction

A ball vanishes from a magician’s closed hand, contradicting what the observer predicted. Surprise marks such moments, when an event departs from current expectations and requires the observer to revise their interpretation of what is happening. Yet how a surprising event reorganizes the geometry of the whole-brain state remains unclear.

Predictive-coding (1–3) and free-energy (4, 5) accounts describe surprise as a signal generated when incoming information deviates from predictions (6), providing a mechanism for updating internal models. However, prediction error primarily characterizes the discrepancy between expected and observed inputs; it does not specify how the entire neural system reorganizes when an event requires a new representation. Natural events rarely violate expectations in isolation: a single surprising event can simultaneously alter sensory interpretation, salience detection, cognitive control, and contextual reconstruction (7, 8). No single brain region captures how these distributed systems interact to generate a new coherent state. Understanding surprise therefore requires a whole-brain framework that characterizes how the configuration of neural states changes when an event exceeds current predictions.

Magic offers a naturalistic paradigm for studying such reorganization. Magic effects disrupt intuitive expectations about physical and causal continuity (9–12), yet these violations occur within coherent event sequences that require observers to reinterpret what has happened. Unlike simplified laboratory violations that isolate individual prediction errors (13, 14), magic events embed unexpected outcomes within meaningful contexts. They therefore capture the representational updating of everyday cognition, in which naturalistic event boundaries can reorganize event-schema representations and reactivate context-specific information from the preceding event (15, 16). Standardized magic-video paradigms allow surprise moments and subjective surprise intensity to be quantified while preserving the richness of real-world event processing (17, 18). Magic is not the object of interest itself, but a controlled naturalistic context for eliciting measurable state transitions.

At the systems level, we hypothesized that the degree of surprise should be reflected in how far the distributed neural state departs from the range occupied during non-surprise viewing. Rather than a change in a single region or a scalar increase in activity, this treats surprise as a displacement of the distributed configuration that represents the ongoing event (19– 21). We refer to such graded reference-relative departures as boundary breakthroughs. The boundary is not a physical border. Theoretically, it corresponds to the range of event states that the current internal model can accommodate; operationally, we estimate it from the range occupied during non-surprise viewing. It is a contour in state space, not a moment in time. A breakthrough occurs when an event carries the neural state past that range.

To make boundary breakthroughs measurable, we represent whole-brain activity patterns in a low-dimensional neural state space, where complex configurations can be characterized by their relative positions and trajectories (22–24). During non-surprise viewing the neural state occupies a range rather than a single point, and the outer edge of that range serves as a reference boundary. A breakthrough is a move beyond it. Boundary exceedance (*E*_BB_, named for that breakthrough) measures how deeply the state moves past the boundary. Because the same video supplies both the reference and the surprise state, the comparison is made within a video rather than across videos. Exceedance is expressed in bits: one bit deeper means the reference distribution reaches that far half as often (Fig. 1a).

**Fig. 1.**
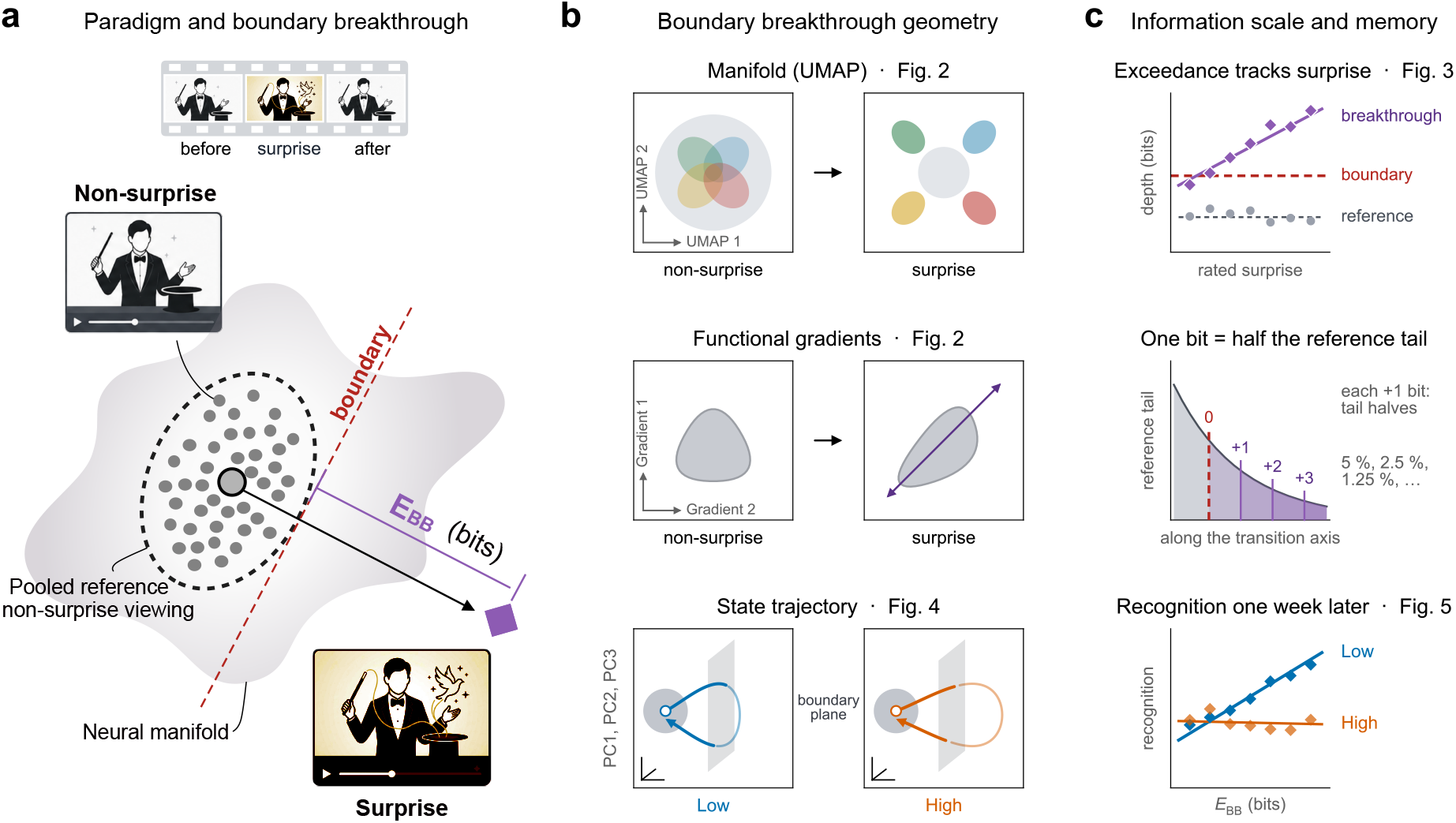
Conceptual and analytic overview of graded whole-brain state departure. **a**, Paradigm and boundary breakthrough. Each magic video runs from an ordinary action (before) through the trick (surprise) to its aftermath (after). An ongoing event is represented by a distributed neural state within a low-dimensional manifold; the pooled non-surprise reference states occupy a region of that manifold rather than a point. A surprise event displaces the state from this reference. The direction of the displacement defines the transition axis, and the outer contour of the reference distribution along that axis defines the boundary. The reference-normalized depth beyond that contour, in bits, is boundary exceedance (*E*_BB_). **b**, Boundary breakthrough geometry, previewed schematically at three levels and developed in Figs. 2 and 4: large-scale networks segregate from an intermixed reference into distinct clusters in a low-dimensional manifold (UMAP); the surprise state expands anisotropically along the functional gradients; and the whole-brain trajectory leaves the reference range, crosses the boundary and returns, further for high than for low surprise. **c**, Information scale and downstream memory, developed in Figs. 3 and 5: exceedance scales with rated surprise in the breakthrough state but not in the reference state; each additional bit halves the reference tail beyond that point; and one week later exceedance is associated with recognition for low-but not high-surprise videos.

We tested one specific prediction: if naturalistic surprise is expressed as a graded departure of the whole-brain neural state beyond its non-surprise reference range, independently rated surprise should scale with the depth of that departure, whereas the paired non-surprise state from the same video should not (Fig. 1). We further asked how this transition unfolds over time and across brain systems (Fig. 1b) and, in an exploratory extension, whether its depth relates to delayed memory (Fig. 1c), without assuming that deeper exceedance is necessarily better or worse. We tested these predictions in the Magic, Memory, and Curiosity (MMC) fMRI dataset (18, 25), in which 49 participants viewed 36 magic videos varying in independently rated surprise intensity.

## Results

The analyses below center on surprise-related events during magic viewing (*Materials and Methods*). We first asked whether surprise reorganized whole-brain state geometry, then tested whether the depth of this departure scaled with independently rated surprise, and then asked how the transition unfolds over time; delayed recognition is treated as an exploratory downstream correlate of boundary exceedance.

### Surprise reorganizes the neural geometry of distributed brain systems

We first asked whether the surprise event displaced the whole-brain neural state relative to the non-surprise reference, that is, whether surprise reorganizes the geometry of the state space. Single-trial whole-brain patterns were estimated for each video event with a GLMsingle-style two-step procedure (26–29), then examined in two low-dimensional views of the same grayordinate data (Fig. 2).

**Fig. 2.**
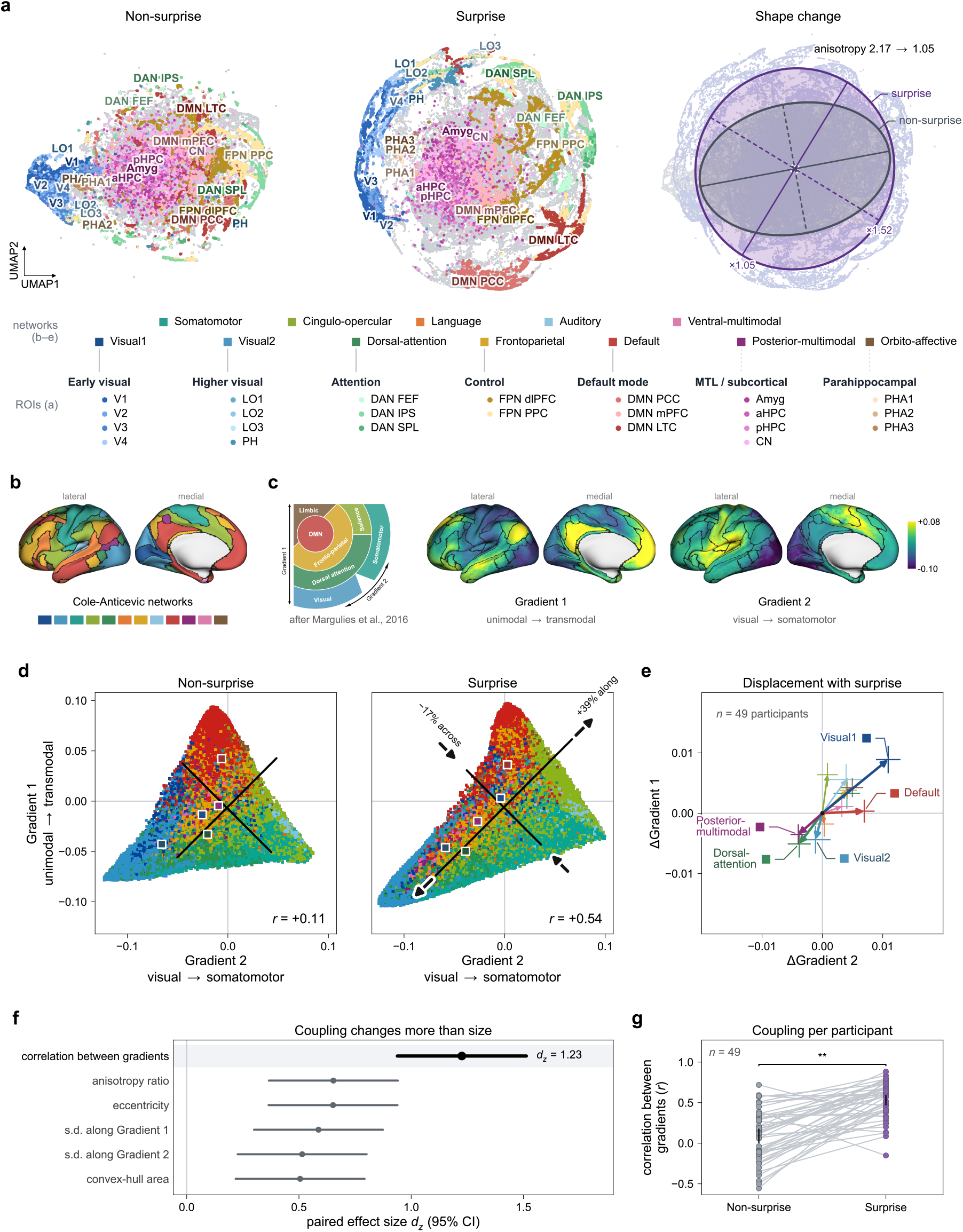
Surprise reorganizes the distributed functional geometry of brain systems. **a**, Whole-brain single-trial activity in a two-dimensional UMAP space fitted jointly on the two states; each point is a grayordinate, colored by region of interest (key below; gray, non-ROI). Right, the two clouds as 2 s.d. ellipses with principal axes: the non-surprise cloud is elongated (anisotropy 2.17) and the surprise cloud nearly isotropic (1.05), scaled 1.05*×* along the long and 1.52*×* along the short non-surprise axis. UMAP distances are not metric. **b**, Cole-Anticevic networks on the inflated left hemisphere (lateral, medial); the strip gives the twelve network colors named in the key of **a. c**, The first two functional gradients of the non-surprise state, organization redrawn after Margulies *et al*. (31). Gradient 1 runs from unimodal to transmodal cortex, Gradient 2 from visual to somatomotor; polarity is fixed by the Procrustes-aligned pre-task resting-state frame. Black lines, network boundaries; one color bar for both maps. **d**, The same gradients as a distribution over the 91,281 grayordinates, colored by network; *r* is the correlation between the gradients. Surprise turns the distribution into a ridge along a common diagonal. Black cross, the two *±* 45^*°*^ directions fixed a priori; its arms are nearly equal under non-surprise and unequal under surprise. Arrows and percentages give the same change estimated within each participant (*n* = 49): +39% along the diagonal, *−* 17% across it. Squares, centroids of the five networks named in **e**; they barely move. **e**, Displacement from non-surprise to surprise: each arrow is one network’s participant-level mean (*n* = 49) with 95% CIs; five are named. **f**, Paired effect sizes (Cohen’s *d*_*z*_ , 95% CI) for six summaries of the distribution, computed per participant (*n* = 49); the change in gradient correlation, shaded, is the largest. **g**, That correlation per participant under the two states (gray lines, one participant; mean *±* 95% CI): 0.09 under non-surprise and 0.53 under surprise, higher in 44 of 49 participants (*t*(48) = 8.58, *p* = 3 *×* 10^*−*11^, *d*_*z*_ = 1.23). Panels **d**–**g** are estimated per participant over grayordinates.

We began by visualizing whole-brain grayordinate activity in a low-dimensional neural manifold with UMAP (30). The embedding separated non-surprise and surprise events (Fig. 2a). In the non-surprise state the grayordinate units formed a relatively compact, intermixed cloud from which primarily visual cortex was differentiated. In the surprise state the major cortical networks pulled apart into distinct clusters. Visual and higher-order visual areas (V1–V4, LO), dorsal-attention (IPS, FEF, SPL), frontoparietal-control (dlPFC, PPC) and default-mode (PCC) systems each formed its own cluster. These clusters remained linked through a shared central core dominated by subcortical structures (hippocampus, amygdala and caudate), so segregation and integration coexisted without the pattern fragmenting (Fig. 2a). Macroscale functional gradients (31–33) provided hierarchical coordinates for characterizing this neural rearrangement (Fig. 2b–g). The surprise state had a larger convex hull, greater dispersion and higher eccentricity than the non-surprise state, an anisotropic expansion rather than uniform diffusion. Estimated per participant over grayordinates, what changed most was not the size of the distribution but its orientation: the correlation between the two gradients rose from 0.09 to 0.53 (Δ = +0.44, 95% CI [0.34, 0.54], *t*(48) = 8.58, *p* = 3 *×* 10^*−*11^, *d*_*z*_ = 1.23), a larger effect than the accompanying growth in hull area, axis-wise spread or eccentricity (all *d*_*z*_ *≤* 0.65; Fig. 2f,g). Which systems moved, and in which direction, was equally systematic (Fig. 2e). Primary visual cortex moved toward the transmodal end of Gradient 1 while dorsal-attention and posterior-multimodal regions moved toward the unimodal end, and both ends shifted on Gradient 2 in the matching direction (SI Appendix, Table S3). The positive shifts in early visual cortex and the negative shifts in association networks therefore form a complementary rather than a uniform pattern, and the two ends move along a common diagonal.

### Exceedance depth scales with rated surprise in the breakthrough state alone

After establishing that surprise reorganized the whole-brain state, we asked whether the depth of that reorganization tracked how surprising an event was. Surprise intensity was indexed by a normative factor combining surprise, interest and curiosity ratings from separate viewers, oriented toward the surprise rating (*Materials and Methods*). We measured each state relative to the distribution occupied during non-surprise viewing, with non-surprise events defined as reference states and surprise events as breakthrough states. In the shared low-dimensional principal component analysis (PCA) space over the 23 regions of interest, the mean reference-to-surprise displacement defined the transition direction (Fig. 3a), and the reference distribution along that direction supplied the boundary against which *E*_BB_ was computed. Most of the axis weight lies on dorsal-attention and frontoparietal cortex (41%) and lateral occipital areas (30%), with 7.5% on early visual cortex; all of these are positive, as are two-thirds of the weights overall. The negative weights fall on medial temporal and default-mode regions (SI Appendix, Table S5).

**Fig. 3.**
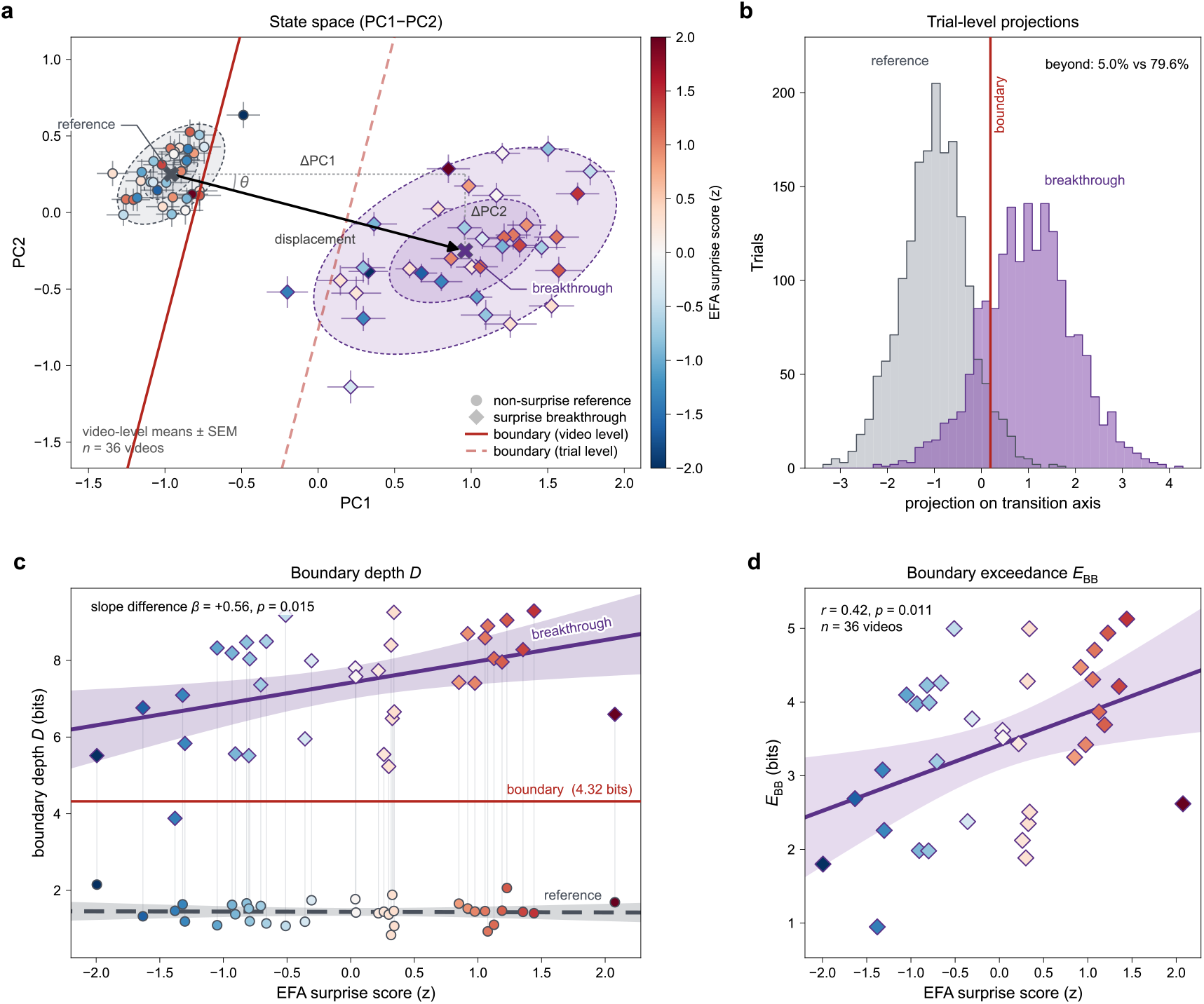
Boundary breakthrough deepens with rated surprise. **a**, State displacement and the reference boundary. Each point is one video’s group-mean pattern in the shared PC1–PC2 space (video-level means *±* s.e.m., *n* = 36 videos), colored by independently rated surprise intensity (EFA surprise score; *Materials and Methods*). Circles, non-surprise reference state; diamonds, surprise breakthrough state; the large gray circle and purple diamond are the centroids and the dashed ellipses 1 and 2 s.d. The black arrow is the mean reference-to-surprise displacement (1.98 in this panel’s units; PC1 and PC2 components dotted), which defines the transition direction. Red lines mark the 95th percentile of the reference distribution along that direction: solid, video-level (the 36 reference video means); dashed, trial-level (the 1,764 reference trials), the boundary used elsewhere in this figure and in Figs. 4 and 5. The direction is estimated from the contrast the panel displays, so the separation shown describes the scale rather than testing it. **b**, How often states lie beyond the boundary. All trials projected onto the transition axis (gray, reference; purple, breakthrough; red line, boundary). By construction 5.0% of reference trials fall beyond the boundary; 79.6% of breakthrough trials do. **c**, Only the breakthrough state scales with rated surprise. Depth is reference-relative, in bits, with the boundary at 4.32 bits. The test uses boundary depth *D* rather than *E*_BB_ so that reference states are not compressed onto the exceedance floor (*Materials and Methods*). Dashed gray fit and circles, reference; solid purple fit and diamonds, breakthrough; 95% confidence bands; thin gray lines connect the two states of each video. The primary test is the paired video-level slope difference, with the video as the only sampling unit (*β* = +0.564 bits per *z*, 95% CI [0.12, 1.01], *t*(34) = 2.56, *p* = 0.015). Points are video-level means of trial-level depth. **d**, Boundary exceedance increases with surprise: video-level *E*_BB_ rose with rated surprise (Pearson *r*(34) = 0.42, *p* = 0.011). Purple line and 95% confidence band; points, video-level means of trial-level *E*_BB_, colored by rating. Convergent with the state-specific scaling in **c**.

On this scale, 79.6% of breakthrough-state trials fell beyond the boundary against 5.0% of reference trials (Fig. 3b). The critical test is whether the depth of exceedance varies systematically with independently rated surprise, whether that relationship is specific to the breakthrough state, and whether it survives estimating the axis and the boundary out of sample. At the video level, the breakthrough state tracked rated surprise intensity but the within-video reference state did not. We tested this dissociation on boundary depth *D*, the same bit scale before the boundary is subtracted. Exceedance is zero for 95% of reference trials by construction, so a comparison on *E*_BB_ itself would rest on a floor rather than a slope (*Materials and Methods*). Because each video contributes exactly one reference and one breakthrough state, the sampling unit is the video, not the individual state. Pairing makes the contrast interpretable: it controls for video-level effects that contribute equally and additively to the two states, but not for effects that differ between states, so the contrast is sensitive to changes at the critical moment rather than to shared video-level characteristics. We therefore regressed each video’s paired breakthrough-minus-reference difference in depth on rating, with the video as the only sampling unit (Fig. 3c). The slope difference was *β* = 0.564 bits per s.d. of rating, 95% CI [0.12, 1.01], *t*(34) = 2.56, *p* = 0.015 (video-cluster bootstrap 95% CI [0.10, 1.00]; paired label-permutation *p* = 0.018). It decomposes into a positive surprise-state slope (*β* = 0.556, *t*(68) = 2.45, *p* = 0.017) and a reference-state slope indistinguishable from zero and tightly bounded (*β* = *−* 0.007, 95% CI [*−* 0.13, 0.12], *t*(68) = *−* 0.12, *p* = 0.90). A state-by-rating model fitted to all 72 state observations gives the same interaction as convergent evidence (*β* = 0.564, *p* = 0.020). In the units of the scale, a video rated one standard deviation above average sat at a breakthrough state about 1.5 times rarer in reference terms.

Exceedance itself scaled with surprise rating across videos (Fig. 3d; Pearson *r*(34) = 0.42, *p* = 0.011; Spearman *ρ* = 0.43, *p* = 0.008; leave-one-video-out *r ∈* [0.37, 0.50], all 36 fits significant). Because this association is confined to the breakthrough state, it can be computed on *E*_BB_ itself, converging with the paired test on the metric’s own scale.

The dissociation also held on the untransformed position along the axis, where both states share one linear scale rather than one that expands differences in the tail (*β* = 0.182 PC units per s.d., 95% CI [0.019, 0.344], *t*(34) = 2.27, *p* = 0.030; reference-state slope *β* = *−*0.003, *p* = 0.91), so it is a property of the geometry itself. More surprising videos were not already in a stronger surprise-related state during their non-surprise segments; they deviated further during the surprise event itself.

Re-estimating the axis and the boundary did not change the result. Refitting both from the other 35 videos for each held-out video gave a primary test of *β* = 0.569 bits per s.d., *p* = 0.015, and refitting the boundary after centering every trial on its own participant’s reference mean left the association intact (SI Appendix, Table S4 and *Construction checks*). Indexing surprise by the single surprise rating rather than the factor score gave the same result (*β* = 0.555 bits per s.d., *t*(34) = 2.47, *p* = 0.019).

### How boundary breakthrough unfolds over time and across systems

Having established that surprise carries the whole-brain state beyond its reference range, we next asked how that transition unfolds over time and across brain systems (Fig. 4). Peri-event responses were estimated with time-resolved finite impulse response models centered on the annotated moment (*Materials and Methods*). Lag 0 is the annotated moment and lags are counted in repetition times (TR = 2 s). We examined these dynamics in two complementary condition-level models: a categorical model comparing the fixed high- and low-surprise video sets, and a separate parametric model testing continuous modulation by independently rated surprise intensity. Because surprise is constant within a video, the high-low contrasts in this section and the next are between two fixed sets of 18 videos.

**Fig. 4.**
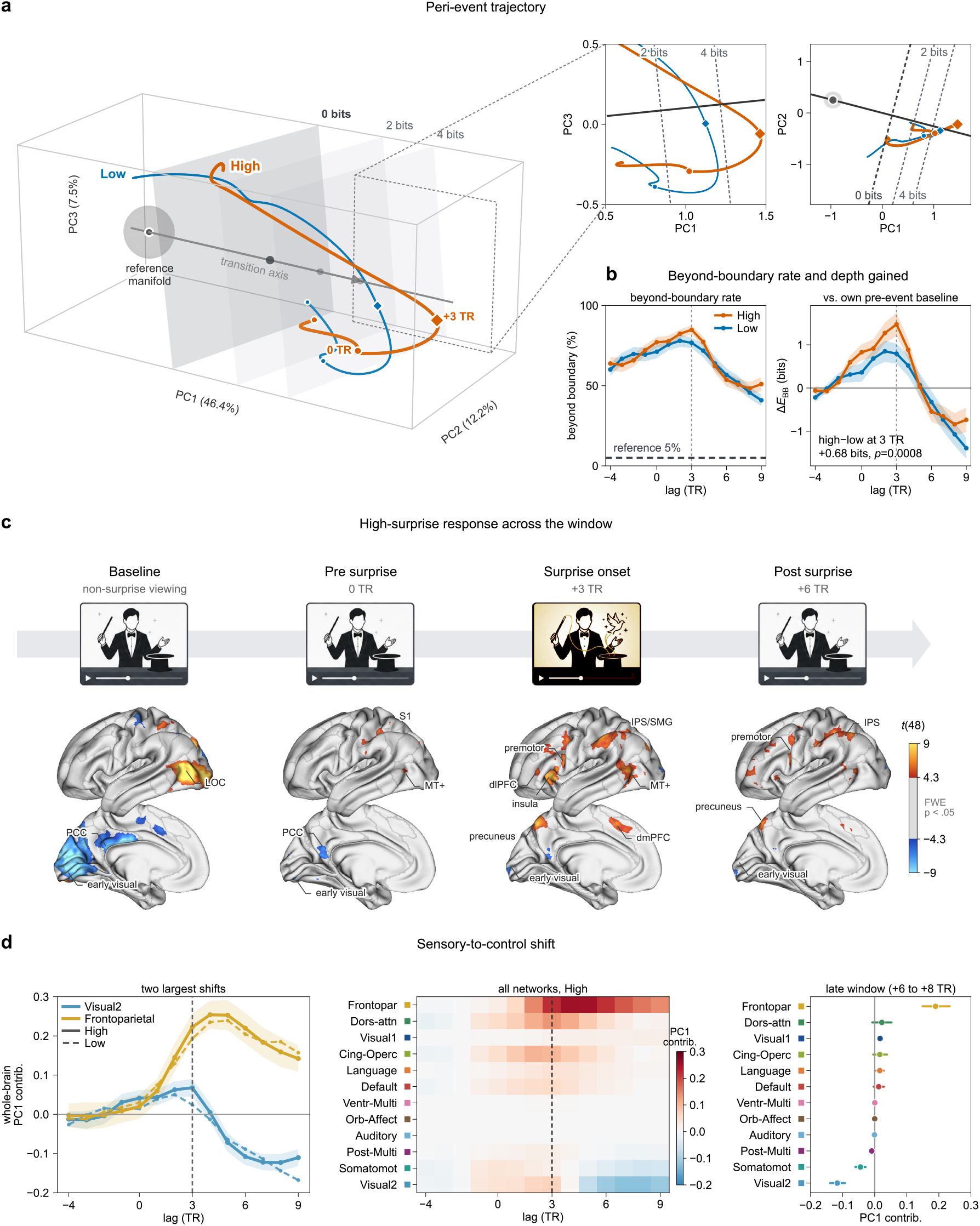
Dynamic realization of boundary breakthrough (*n* = 49; lag 0 is the annotated moment; vermilion, high surprise; blue, low surprise). **a**, High- and low-surprise events leave the reference region along different paths. Peri-event trajectories are projected into the frozen trial-level PCA of Fig. 3 (marker shapes as there). The axis runs from the reference centroid to the surprise centroid; three planes of constant exceedance cross it at 0, 2 and 4 bits, the darkest being the boundary, *E*_BB_ = 0. Planes and panel **b** use the trial-level reference; gray sphere, 1 s.d. of the 36 video means, inside the boundary. Right, the same trajectories on the PC1–PC3 and PC1–PC2 planes; dashed lines, the 0-, 2- and 4-bit traces. **b**, The event deepens an excursion already under way. Left, proportion of trials beyond the boundary at each lag; dashed line, the 5% of reference trials beyond it by construction. Right, depth gained relative to each trial’s own pre-event baseline, Δ*E*_BB_; at +3 TR the high-surprise set had gained +0.68 bits more than the low-surprise set (*t*(48) = 3.56, *p* = 0.0008). Points, one per TR; shading, 95% CI across participants. **c**, Time-lagged GLM maps for high-surprise videos at 0, +3 and +6 TR after the annotated moment (frames above sketch the trick), next to the non-surprise reference term. Left hemisphere, lateral above and medial below; *t*(48), TFCE-FWE *p <* 0.05; gray outlines, Cole-Anticevic networks; warm, positive; cool, negative *t* (not the low-surprise condition). Both hemispheres, all windows and the low-surprise and parametric models: SI Appendix, Fig. S1. **d**, A sensory-to-control redistribution, clearest late in the response. Left, the two networks with the largest pre-event-to-late change: frontoparietal stays above its pre-event baseline, Visual2 falls below it (solid, high surprise, bootstrap 95% CI; dashed, low surprise). Middle, all twelve networks under high surprise, network by lag, same scale. Right, late-window (lag 6–8) contribution of every network (bootstrap 95% CIs). Contributions are to PC1 of the whole-brain decomposition and support the ordering of networks, not absolute values.

We began with the state itself, in the coordinates Fig. 3 established. Peri-event activity was projected into the same frozen trial-level principal component analysis that defines the boundary geometry there, so panels a and b share one coordinate system and one non-surprise reference with Fig. 3 (*Materials and Methods*). High- and low-surprise events left the reference region along different paths, the high-surprise trajectory reaching the more extreme position along the transition axis (Fig. 4a).

A trajectory shows where the state went; exceedance converts that into how far outside the reference range it is, lag by lag (Fig. 4b). The proportion of trials beyond the boundary was highest in the response windows following the annotated moment (85% high and 77% low at +3 TR; Fig. 4b, left), and was already 64% and 60% at the start of the modeled window (lag *−*4), so the peri-event excursion rises from a raised floor rather than from the 5.0% of reference trials that lie beyond the boundary by construction. The depth gained beyond the boundary, Δ*E*_BB_, peaked in the same windows, and at +3 TR the 18 high-surprise videos had gained more depth than the 18 low-surprise ones (+1.474 versus +0.795 bits, each referenced to the trial’s own pre-event baseline). The +0.68-bit difference is reliable across participants (*t*(48) = 3.56, *p* = 0.0008; Fig. 4b, right), who all viewed the same two sets.

The two conditions had already begun to diverge before the annotated moment. Across lags, the high-minus-low difference in state displacement along the same axis formed three familywise-error-corrected temporal clusters; the earliest, at lags *−*1 to 0 (*p* = 0.044), begins before the annotated moment and therefore cannot be a hemodynamic response to it, and by that point both conditions are already above their pre-event baseline (SI Appendix, Fig. S4). What the event does is deepen an excursion that is already separating the two conditions. Anticipation, imprecision in the annotated moment and slow build-up within the trick cannot be separated here.

Having located when the state ran deepest, we asked which systems carried it there. The answer was a redistribution rather than a uniform increase in activity. Non-surprise segments provided the viewing-state reference against which depth and exceedance were defined; relative to it, the high-surprise response recruited visual, parietal and control-related regions, sparse before the annotated moment, most extensive at the +3-TR response window and receding by +6 TR (Fig. 4c). In the direct high-minus-low contrast the only corrected clusters after the annotated moment were visual, all in the low-greater-than-high direction (one pre-event cluster in left 7Am ran the other way; SI Appendix, Table S1), whereas the parametric model showed positive modulation by rated surprise across visual, dorsal-attention and control regions. The two models answer different questions. The categorical contrast compares the mean responses of two fixed sets of 18 videos, whereas the parametric term estimates, across all 36 videos and after the shared event response is accounted for, whether the response scales with the continuous rating. The same visual regions can therefore show a larger mean response in the low-surprise set and positive modulation by rating. Condition-specific maps showed prominent frontoparietal and control-related responses in the high-surprise set, including right anterior insula, left anterior intraparietal cortex (AIP), angular gyrus and bilateral parietal cortex, whereas the low-surprise set showed stronger and more sustained visual responses, including ventromedial visual cortex (VMV3), V4 and early visual cortex. Peak statistics for all four peri-event windows are given in the SI Appendix, with the corresponding maps and subcortical effects in Figs. S1 and S2.

At the network level this sensory-to-control redistribution became clearest after the event (Fig. 4d). Of the twelve Cole-Anticevic networks, the frontoparietal network and higher visual cortex (Visual2) showed the most extreme post-event values. Both peaked positive in the response windows following the annotated moment (Visual2 at lag +2, frontoparietal at lag +5), so the peak does not separate them; they diverge afterwards, and in sign. Over lags +6 to +8, after the shared early response has resolved, the frontoparietal contribution stayed above its pre-event baseline (0.188, bootstrap 95% CI [0.149, 0.226]) whereas Visual2 fell below it (*−* 0.118, 95% CI [*−* 0.139, *−* 0.097]). All twelve networks are shown without selection in Fig. 4d, middle and right, with both conditions’ full time courses in the SI Appendix (Fig. S3). At the finer parcel scale the two conditions separated by sign rather than by magnitude (SI Appendix, Fig. S5).

### Exceedance depth relates to later recognition memory

Finally, boundary exceedance was mirrored in behavior one week later. Participants recognized the 18 low-surprise videos more accurately than the 18 high-surprise ones (0.661 versus 0.600 against a chance level of 0.25; paired *t*(48) = *−* 3.13, *p* = 0.003; Fig. 5c), with no reaction-time difference (*p* = 0.848). Across the same participants, those 18 high-surprise videos also pushed the state further past the boundary (3.15 versus 3.68 bits, +0.52 s.d., *t*(48) = 4.69, *p <* 0.0001; Fig. 5b). The two effects therefore run in opposite directions: deeper exceedance, poorer memory.

**Fig. 5.**
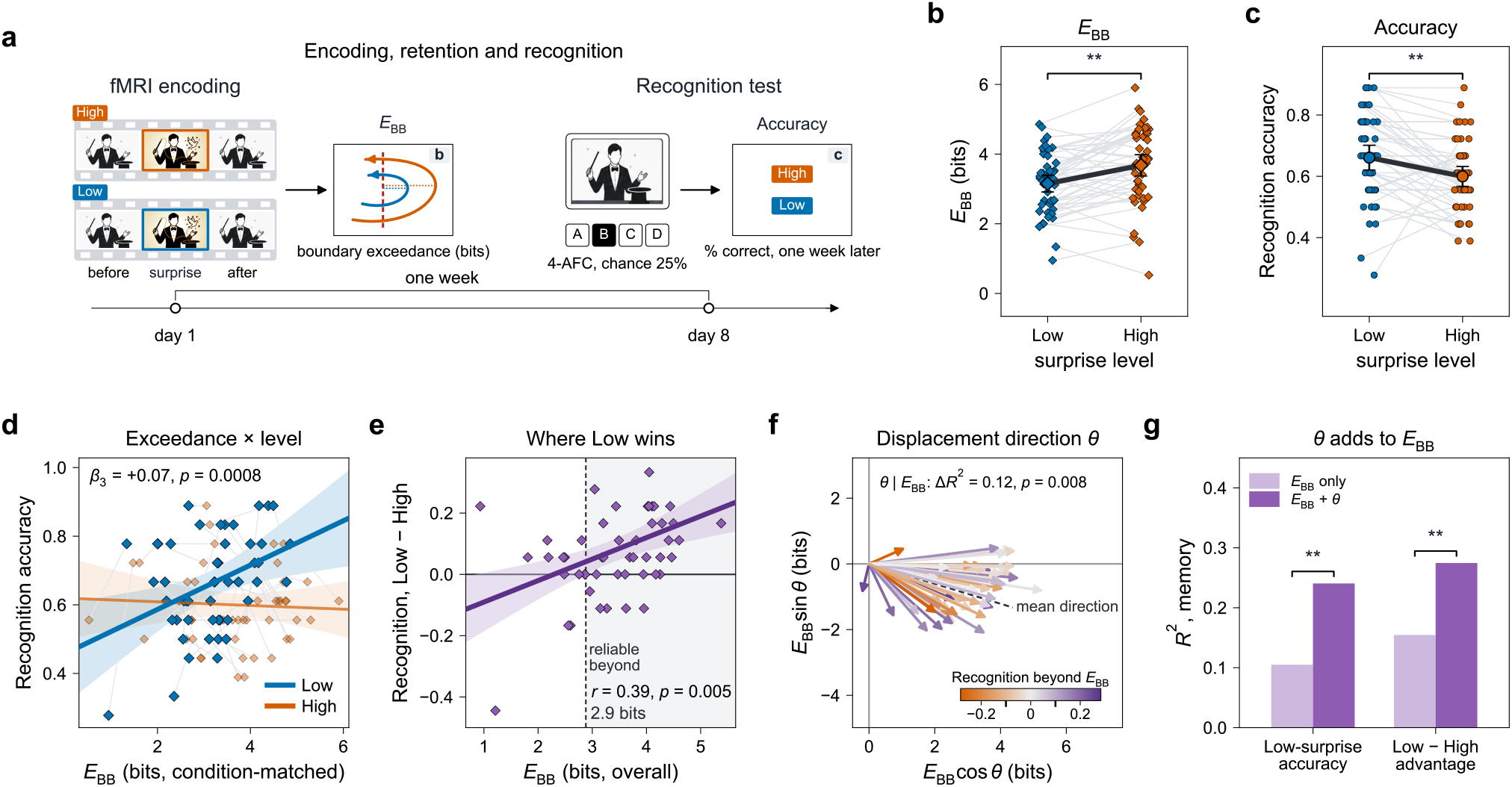
Boundary exceedance and one-week recognition memory. Participant-level analyses (*n* = 49); *E*_BB_ is the exceedance beyond the boundary of Fig. 3, computed per trial and averaged within participant and condition. Vermilion, high surprise; blue, low surprise. **a**, Design. Each of 36 magic videos (18 per level) was viewed during fMRI on day 1; the whole-brain state during viewing gives one *E*_BB_ per video: a trajectory leaves the reference region, crosses the boundary (dashed red) and returns, further for high surprise. One week later a still image from each video cued a choice among four verbal descriptions of the trick (chance 25%). Small letters mark the panels showing each quantity. **b**, High-surprise videos push the state further past the boundary: 3.15 versus 3.68 bits, paired *t*(48) = 4.69, *p <* 0.0001. Thin lines, participants; large markers, means *±* 95% CI. **c**, Yet they are remembered worse: 0.661 versus 0.600, *t*(48) = *−*3.13, *p* = 0.003. **d**, Exceedance is associated with memory only for low-surprise videos. Each participant contributes one point per condition, joined by a gray line; fits with 95% confidence bands span the common *x* range. Standardized *E*_BB_ *×* surprise interaction *β*_3_ = +0.071, 95% CI [0.032, 0.110], *t*(53.7) = 3.54, *p* = 0.00085 (participant random-intercept model, Satterthwaite d.f.). **e**, Model-implied low-over-high recognition advantage as a function of each participant’s overall *E*_BB_ (line, *±* 95% CI); the advantage is reliable beyond 2.9 bits (shaded); observed association *r* = 0.39, *p* = 0.005. Overall *E*_BB_ averages trials inside and beyond the boundary, and the threshold is specific to this sample. **f**, Each arrow combines one participant’s mean exceedance *E*_BB_ as its length with the angle *θ* of their mean displacement in the PC1–PC2 plane; it is not the untransformed displacement vector. Dashed line, circular mean. Arrows are colored by low-surprise recognition after regressing out *E*_BB_ (purple, better than expected; vermilion, worse). Adding *θ* to a model with *E*_BB_ improves prediction of the low-over-high advantage (Δ*R*^2^ = 0.12, *p* = 0.008). **g**, The same gain for both outcomes: variance explained by *E*_BB_ alone and with *θ* (low-surprise accuracy, 0.105 *→*0.241, *p* = 0.006; low-over-high advantage, 0.155 *→*0.275, *p* = 0.008; *F* tests for the added term). *θ* alone is unrelated to memory.

In an exploratory analysis, we then asked whether the exceedance–memory relation depends on surprise level. Across participants, who are the sampling unit here rather than videos, we modeled recognition on exceedance, surprise and their interaction (*Materials and Methods*). The mnemonic benefit of deeper exceedance was present in the low-but not the high-surprise condition (Fig. 5d). The standardized *E*_BB_ *×* surprise interaction was *β*_3_ = +0.071, 95% CI [0.032, 0.110], *t*(53.7) = 3.54, *p* = 0.00085, from a participant random-intercept model with Satterthwaite denominator degrees of freedom. Consistent with this moderation, greater exceedance was associated with better recognition for low-surprise videos (*β* = 0.065, *t*(47) = 2.87, *p* = 0.006) but not for high-surprise videos (*p* = 0.729). It was also associated with a greater recognition advantage for low-over high-surprise videos (*β* = 0.060, *t*(47) = 2.93, *p* = 0.005). With the boundary re-estimated from the other 48 participants and applied to each held-out participant, that association was *r* = 0.39, *p* = 0.005.

The bit scale lets this last relation be located on an interpretable, reference-relative axis. In the present sample the model-implied low-over-high advantage became reliable beyond approximately 2.9 bits past the boundary (Fig. 5e).

Because the displacement is a vector, we also asked, in an exploratory analysis, whether its direction carried information beyond how far it reached past the boundary. Adding the displacement angle to a model already containing *E*_BB_ improved prediction of the low-over-high memory advantage (Δ*R*^2^ = 0.120, *F* (1, 46) = 7.61, *p* = 0.008; leave-one-out cross-validated *r* rose from 0.22 to 0.31; Fig. 5f,g), and of low-surprise recognition itself (Δ*R*^2^ = 0.136, *p* = 0.006; Fig. 5g). The angle alone was unrelated to memory, a direction-given-depth effect. This geometry–memory relation was robust to trait and state curiosity and to need-for-cognition. The full memory models and the direction and curiosity analyses are given in the SI Appendix.

## Discussion

### Surprise as a graded boundary breakthrough of the neural state

The central finding is a graded relationship between surprise intensity and whole-brain state departure. Across videos, independently rated surprise tracked the depth of the breakthrough state but not the paired within-video reference state, in the paired video-level test and in the rating *×* state model, and the relationship persisted when the axis and the boundary were estimated from the other videos and applied to each held-out video. The ratings are normative ones collected from separate samples of viewers (17, 34), so the geometry predicts how a different group of people experienced the same videos, not what these participants reported. The scaling is also confined to one of the two states within each video; this state specificity argues against effects shared equally by the two states.

The graded relationship links the intensity of surprise to the position of a distributed neural state relative to an observed reference range. Rather than contrasting surprising with unsurprising events, or relating surprise to the amplitude of responses in particular regions, the result maps the degree of surprise onto how far the whole configuration lies beyond the pooled reference range observed during non-surprise viewing: a video rated one standard deviation above average sat at a state about 1.5 times rarer in reference terms. The graded association is consistent with departures of varying depth, although whether these reflect continuous changes in what the current event model can accommodate remains to be tested; the observed boundary is an operational reference, not a direct measure of that model. Surprise can therefore be expressed as a measurable, reference-relative neural geometric quantity (Fig. 3).

### Whole-brain reconfiguration underlying boundary breakthrough

Surprise did not translate the whole pattern; it changed its shape. In the macroscale gradient space the surprise state had a larger convex hull, greater dispersion and higher eccentricity, and its ends moved in opposed directions. Early visual cortex shifted positively along the first two gradients while dorsal-attention and posterior-multimodal systems shifted negatively (Fig. 2e). Neither global nor region-specific positive gain rescaling can produce this reconfiguration: scaling a region’s signal leaves its Pearson correlations, and therefore the gradient geometry, unchanged.

Because sensory and association systems move against each other along a common diagonal, two nearly independent gradients become coupled. The gradient analysis characterizes changes in macroscale functional organization. In a separate region-of-interest activity space, the mean reference-to-surprise displacement defines the direction along which we test the relationship between surprise intensity and state departure (Fig. 3); the two analyses provide complementary descriptions of surprise-related reorganization rather than one mapping onto the other. The UMAP embedding shows the same reorganization at the level of grayordinates, with the major networks separating into distinct clusters that stay linked through a shared subcortical core, so segregation and integration coexist rather than trade off.

### From sensory processing to control: what the time course implies

Boundary breakthrough unfolds as a process rather than a moment. Cognition draws on the dynamic integration of activity across systems (35), and a breakthrough is one such reorganization. Both conditions reached their deepest excursion in the same response window to within one lag, so what separates them is distance (Fig. 4b); the pre-onset divergence is the one timing difference the data show. That distance is the component along the transition axis, with Δ*E*_BB_ reading each trial against its own pre-event state, so the conditions differ in how unusual their configuration becomes in the reference distribution’s own terms, a question raw response amplitude cannot answer.

The whole-brain maps and the region-of-interest time courses converged on the same asymmetry (Fig. 4c): the conditions were distinguished by a redistribution of activity between visual-parahippocampal and default/control systems rather than by a uniform difference in amplitude. A separate parametric model identified rating-related modulation across visual, dorsal-attention and control regions. Low-surprise events appear to evoke sustained adjustment of visual detail and context, whereas high-surprise events recruit systems for monitoring, attentional selection and updating of the current internal model, consistent with high-level models constraining sensory interpretation (5) and with the sensory-to-transmodal hierarchy (31). What the time course adds is when this happens: the redistribution is clearest after the shared early response has resolved, with the frontoparietal contribution staying above its pre-event baseline while higher visual cortex falls below it (Fig. 4d). The shift converges with cross-context predictive work in which belief-inconsistent surprise is carried by frontoparietal, medial-frontal and default-mode interactions and lower surprise by visual and motor ones (7).

The state was already moving before the annotated moment, and recent intracranial evidence offers a candidate mechanism: human hippocampal ripples increase before stimulus presentation under higher predicted uncertainty and modulate the subsequent visual cortical response to surprising input (36). fMRI cannot resolve ripple-scale mechanisms, but the data fit the broader principle that natural surprise is a state transition shaped by what the observer already expected.

### Boundary exceedance, prediction error and event segmentation

Predictive-coding research asks how error signals arise and propagate; event-cognition research asks how boundaries segment continuous experience. Boundary exceedance provides a testable state-space description linking the two, because its boundary is instantiated rather than asserted: the operational boundary is a quantile of the observed reference distribution, and exceedance measures how far the state departs from the range that non-surprise viewing occupies across the task, an observable proxy for an event exceeding what the current model can accommodate. *E*_BB_ differs from prediction error because it describes where the distributed neural state lies relative to its empirical reference range, whereas prediction error characterizes the mismatch signals that violated expectations generate, signals that bias hippocampal states and disrupt existing representations as episodic models are updated (37, 38). The two are complementary rather than one a relabelling of the other.

Exceedance describes position relative to a reference range, whereas event segmentation concerns changes in the organization of ongoing experience (39, 40). Segmentation asks whether the state has changed; exceedance asks how far outside the reference range it has moved, on a stated scale. Their temporal correspondence remains an empirical question, testable by comparing exceedance trajectories with boundaries recovered by a hidden-state event model on the same data.

The ingredients of the scale are established, Shannon self-information (41) applied to empirical tail probabilities (42– 44), and two properties matter for its interpretation. The transition direction distinguishes breakthrough from undirected outlyingness, which would count a departure in any direction, including those orthogonal to the one the event produces. And the resulting bit scale is reference-relative: what transfers across datasets is its probabilistic interpretation, not the numerical value itself (invariance and sensitivity analyses in the SI Appendix). Because a natural event mixes sensory change, causal violation, attentional reallocation and contextual integration at once, no single region, rating or time point characterizes it; exceedance offers a testable systems-level intermediate variable for that composite event.

### When boundary exceedance supports cognition

Boundary exceedance was mirrored in behavior in a way that argues against a simple rule in which deeper exceedance always aids memory. The low-surprise set was better recognized than the high-surprise set, and across participants deeper exceedance was associated with better recognition only in the low-surprise condition (Fig. 5). This fits event-boundary work in which boundaries aid encoding but excessive discontinuity impairs integration across a continuous context (21, 45, 46). In these same videos, encoding is enhanced when viewers are curious about the outcome (47); the condition-dependent association observed here motivates the hypothesis that the mnemonic consequence depends less on how far the state moves than on whether the resulting transition can still be integrated with the current event model (48, 49). This integrability account is a hypothesis to be tested directly, not a quantity measured here.

What depth cannot capture is the direction of the transition: exceedance is a projection onto the transition axis, so the orthogonal component of the displacement is discarded by construction. The Results treat that component as a direction-given-depth effect: the angle alone was unrelated to recognition, but added to a model already containing *E*_BB_ it improved prediction. Depth and direction may therefore carry dissociable behavioral information. Testing this would require a direction measure defined in the full state space and a prespecified hypothesis in an independent sample.

### Limitations and future directions

The main limitation is that all analyses derive from a single dataset. Leave-one-video-out and participant-level cross-validation establish internal robustness, but not dataset-level generalization; the stability of the displacement and exceedance results should be tested in independent samples and with larger video sets. The geometry rests on low-dimensional approximations (PCA, UMAP and functional gradients) whose convergence supports the state-space results without establishing a unique neural manifold. Exceedance depends on the selected state space, transition axis and reference distribution (SI Appendix, Table S4). The reference distribution was pooled across video-onset states and therefore represents task-wide non-surprise viewing rather than the predictive range of the current event. Differences in temporal position may also contribute to the contrast between reference and surprise states. The naturalistic design brings its own limits. Magic videos place prediction violation in a context where visual salience, causal violation, curiosity and contextual continuity are not fully separable. Curiosity is the clearest case: it modulates episodic memory (50, 51) and covaries with surprise in these materials (47), and although the geometry–memory relation survived participant-level curiosity controls, its behavioral relevance cannot yet be uniquely attributed to surprise. The temporal resolution of fMRI also limits what the peri-event sequence can reveal; EEG, MEG or intracranial recordings would be needed to resolve that sequence, and experimental manipulation of boundary strength to test the functional meaning of exceedance across paradigms.

## Conclusion

In the human brain, naturalistic surprise is expressed as a graded boundary breakthrough: more surprising events carry the whole-brain state farther beyond a measured non-surprise reference range. The scale is defined by a reference distribution, a transition axis and a chosen tail probability; the reference range is estimated from the data rather than assumed. Boundary exceedance places the departure on a reference-relative information scale, but the empirical result is the state-specific scaling of breakthrough depth with independently rated surprise.

## Materials and Methods

Detailed methodological information is provided in the Supplementary Materials. In brief, we analyzed the publicly available MMC fMRI dataset (18, 25). That dataset was collected under a protocol approved by the University Research Ethics Committee of the University of Reading (UREC 18/18), and all participants gave informed written consent (18). No new data were collected for the present study, and no separate ethical approval was required. Of the 50 healthy right-handed adults in the original dataset, one was excluded because of incomplete scanning data, leaving 49 for analysis. The original MMC dataset description gives the data-collection workflow and dataset structure. Before scanning, participants completed online questionnaires. During the same fMRI session, each participant completed pre-task resting-state BOLD scanning, magic-viewing task scanning and post-task resting-state BOLD scanning. The resting-state runs enter the present analyses only as a common reference frame for the gradient geometry. The task included three functional runs in which participants watched 36 magic videos. Each video contained one or more expert-annotated objective surprise moments. Video surprise intensity was the *z*-standardized score from a one-factor exploratory factor analysis of surprise, interest and curiosity. These three were retained from five normative ratings (clarity, surprise, interest, solution confidence and curiosity) by their association with the surprise anchor (|*r*| *>* 0.80, *p <* 0.05), and the factor was oriented so that higher scores indicate greater surprise. Videos were divided into high- and low-surprise sets on this score. One week later, participants completed a delayed memory test in which cued recall preceded recognition. Recognition was four-alternative forced choice: a cue image from each of the 36 videos appeared with four verbal descriptions of what happens in that trick, one of them correct, so chance was 25%. The measure is therefore memory for a trick’s content rather than familiarity with the clip. The primary behavioral measure was delayed recognition accuracy, and recognition reaction time was used in supplementary analyses.

Anatomical and functional images were preprocessed with fMRIPrep 25.0.0 (52), whose surface reconstruction uses FreeSurfer (53), and resampled to fsLR grayordinate space (54). Two estimates of the event response were then derived from these data, a single-trial estimate and a time-lagged estimate, and each analysis below uses whichever one its question requires.

The single-trial estimate defines the state geometry in which exceedance is measured (Figs. 2, 3 and 5). Single-trial activity patterns were estimated with a two-step procedure following the HRF-selection and noise-regression stages of GLMsingle (26), without its fractional-ridge shrinkage step. Each pattern was reduced to one value per region of interest, the mean over that region’s grayordinates, where the regions are the 23 a priori masks whose intersections with the HCP-MMP and Cole-Anticevic base parcels (55, 56) give the 162 parcel units used for the parcel-resolved analysis. Principal component analysis over those 23 values gave the PC1–PC2 plane. Every quantity defined below, including the displacement, the transition axis, the reference distribution and boundary exceedance, lives in that plane rather than in the trajectory space of the time-lagged estimate. A uniform manifold approximation and projection (UMAP) embedding of all cortical and subcortical grayordinates provides visualization only, because its axes are not metric. BrainSpace macroscale functional gradients aligned to the pre-task resting-state frame provide a separate set of metric hierarchical coordinates. These were estimated per participant over grayordinates, which is what Fig. 2 displays and which allows the geometric summaries to be tested across participants. PCA and UMAP were always fitted jointly on the states being compared, so that non-surprise and surprise observations occupy a single common embedding rather than separately fitted spaces.

The time-lagged estimate characterizes when and where the response unfolds (Fig. 4). Three finite impulse response (FIR) models were fitted, none of which assumes a canonical hemodynamic response. Two condition-level FIR general linear models (GLMs) centered on the annotated surprise moment gave the whole-brain maps of Fig. 4c: one estimated the boundary-event response together with continuous parametric modulation by rated surprise, and a separate model estimated the high- and low-surprise conditions and their direct contrast. Group-level inference used PALM permutation testing and threshold-free cluster enhancement (TFCE-FWE) (57, 58). A video-wise least-squares-separate FIR model (27) gave one coefficient per video and lag; region-of-interest time series extracted from it in the HCP-MMP and Cole-Anticevic parcellation supplied the peri-event trajectory and exceedance time course (Fig. 4a and b, by principal component analysis over the 23 target regions) and the network contributions (Fig. 4d, from a whole-brain decomposition). Lag 0 is the annotated moment and lags are counted in repetition times (TR = 2 s). The hemodynamic response is slow enough that adjacent lags sample overlapping activity, so a lag labels a blood-oxygen-level-dependent (BOLD) sampling window rather than a temporally resolved neural stage.

Boundary exceedance (*E*_BB_) was the operational measure of boundary breakthrough. Within each video, the displacement runs from its non-surprise reference state to its surprise state; the transition axis is the group-level mean of those displacements, normalized to unit length. Pooling non-surprise trials across videos and participants then yielded the task-wide reference distribution, whose upper-tail contour along that axis defined the operational boundary. Note that the displacement is a within-video contrast whereas the reference distribution is pooled, so exceedance is normalized against non-surprise viewing across the task. For a state *x*, the empirical upper-tail probability *π*(*x*) is the proportion of reference states whose projection onto the transition axis is at least as far out as that of *x*. The direction-specific boundary was the contour *π*(*x*) = *q*, with *q* = 0.05. Following the Shannon self-information transform (41, 42), we expressed a state’s boundary depth as

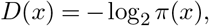

and introduced boundary exceedance as

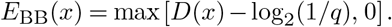

Thus the boundary is fixed at log_2_(1*/q*) = 4.32 bits; *E*_BB_ = 0 for states on or inside the contour; and each additional bit beyond it halves the reference probability of reaching at least that far. Accordingly, *E*_BB_ serves as a probability-calibrated observational scale. Within a specified reference model, equal one-bit increments carry the same reference-relative interpretation, even when the raw state-space coordinates are hard to interpret.

Because *E*_BB_ is zero for every state on or inside the boundary, tests that require variation within the reference distribution were computed on *D*; these are the paired rating test and the state-by-rating model. *E*_BB_ was used wherever the quantity of interest is exceedance itself. Above the boundary, *E*_BB_ equals *D* minus the constant boundary depth of 4.32 bits; on or inside the boundary, *E*_BB_ is set to zero whereas *D* retains within-boundary variation.

For the peri-event analyses, exceedance was also expressed relative to each trial’s own pre-event state: Δ*E*_BB_ is the lag-wise value minus the mean *E*_BB_ over lags *−*4 to *−*2. The correction is needed because the pre-event state is already displaced (64% and 60% of trials beyond the boundary at lag *−* 4), so an absolute value would mix event-driven movement with pre-existing position. Because *E*_BB_ is a log tail probability, the difference of two values is the log_2_ ratio of reference tail probabilities before and after the event, against the same reference. The floor at zero makes that reading approximate for the 17% of trial baselines that lie inside the boundary; the same contrast on the uncensored depth *D* is larger (+2.00 versus +1.08 bits), so the censoring is conservative. The reference distribution was defined at the trial level (*n* = 1,764 non-surprise trials); video- and participant-level values are averages of trial-level *D* or *E*_BB_. Let *k*(*x*) be the number of reference trials at or beyond *x* along the transition axis. We used the finite-sample estimate 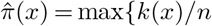, 1*/*(*n* + 1) , so that *D* is bounded above at log_2_(*n* + 1) = 10.79 bits and no state receives an infinite score. Because the reference distribution is empirical, the scale is censored at that ceiling, which 23.0% of breakthrough trials reach. By collapsing the most extreme observations, this censoring is expected to attenuate rather than amplify monotonic surprise scaling. Rank-based, censored-regression and parametric-tail sensitivity analyses (SI Appendix) confirm that the reported association does not depend on ties at the top of the empirical distribution.

We cross-fitted the axis and boundary within the shared state-space basis to test whether surprise scaling depended on estimating them from the same videos. For each held-out video, the transition axis and the trial-level reference distribution were estimated from the other 35 videos and then applied to that video. Label permutation repeated this full axis-and-boundary estimation pipeline. We additionally repeated the analysis after centering each trial on its participant-specific reference mean. For the geometry–memory relation, the boundary was estimated from the other 48 participants and applied to each held-out participant.

The primary test of state-specific surprise scaling was performed at the level of the video. Because each video contributes exactly one reference state and one breakthrough state, the video is the sampling unit. We therefore regressed the paired difference Δ*D*_*j*_ = *D*_breakthrough,*j*_ *− D*_reference,*j*_ on rating across the 36 videos (df = 34). A video-cluster bootstrap and a paired label permutation provided distribution-free checks. The state-by-rating model fitted to the 72 state observations is reported as convergent evidence. It uses HC3 robust standard errors, because residual variance differs markedly between the two states, with *t*-distribution inference on its 68 residual degrees of freedom. Behavioral and delayed-recognition relationships were analyzed with paired *t* tests and mixed-effects regression.

## Supporting information

Supplementary Information (Supplementary Methods, Results, Tables 1-5, Figs. 1-5)

## Data and code availability

The MMC dataset (18) is available from OpenNeuro (ds004182, version 1.0.1, https://doi.org/10.18112/openneuro.ds004182.v1.0.1) (25). The normative stimulus ratings used to index surprise intensity are those distributed with the MagicCATs collection (17), available from the Open Science Framework (https://osf.io/ad6uc/). The videos themselves are distributed by the MagicCATs authors, who release them on request for research use. A reference implementation of boundary exceedance is available at https://github.com/ibrainlab/boundary-exceedance. It covers the estimation of the transition axis, the pooled reference distribution, the empirical tail probability and its bit transform. Main software packages included fMRIPrep, FreeSurfer, FSL, ANTs, Connectome Workbench, PALM, GLMsingle, scikit-learn, umap-learn and BrainSpace.

## ACKNOWLEDGEMENTS

We thank the creators of the MMC dataset and of the MagicCATs stimulus collection for making these resources available for research. We gratefully acknowledge the developers of fMRIPrep, FreeSurfer, FSL, ANTs, PALM, BrainSpace, GLMsingle and the Connectome Workbench for providing open-source neuroimaging tools that made this work possible. This work was supported by the National Natural Science Foundation of China (grant no. 32300876) and by the Beijing-Hetian Joint University Laboratory Construction Project.

## AUTHOR CONTRIBUTIONS

Y.H. and C.J. designed research; Y.H. and C.J. performed research; Y.H. and C.J. contributed new analytic tools; Y.H., J.L. and C.J. analyzed data; Y.H., J.L. and C.J. prepared the figures; Y.H., J.L. and C.J. wrote and revised the paper; and C.J. supervised the project.

