## Supplementary Information (Supplementary Methods, Results, Tables 1-5, Figs. 1-5) for "The neural geometry of naturalistic surprise: Graded boundary breakthroughs in the human brain"

### Supplementary Information for The neural geometry of naturalistic surprise: Graded boundary breakthroughs in the human brain

**Yunqiang Hao**[1] **and Chao Jiang**[1,✉]

[1]School of Psychology, Capital Normal University, Beijing 100048, China

#### Supporting Information Text

#### Supplementary Methods

##### Participants, stimuli and behavior

The MMC dataset (OpenNeuro ds004182) (1) contains 50 adults. One was excluded for incomplete scans, leaving 49 (35 female; 18–37 years, mean 25.4, SD 5.2). The original control-versus-reward grouping was not used. Each of the 36 magic videos has one or more expert-annotated moments at which the effect becomes visible. When several fell within 8 s, the first was the primary event and the rest were modeled as events of no interest. Surprise intensity is the $z$-scored one-factor score of the normative surprise, interest and curiosity ratings (2, 3), the three of five ratings correlated with surprise at $|r| > 0.8$. The factor was oriented positively with surprise, and a median split gave 18 high- and 18 low-surprise videos, held constant across all analyses. Delayed recognition was four-alternative forced choice on a still cue image (chance 25%). All 36 studied videos were tested and no unstudied videos were included; the construction of the incorrect options is not reported in the dataset publication. Accuracy was coded as binary per participant and video. Reaction time was converted to seconds, with non-positive values treated as missing. Recall and confidence ratings are not analyzed. Trait Curiosity, State Curiosity and Need for Cognition serve as covariates below.

##### Acquisition, preprocessing and regions of interest

Functional images were single-band gradient-echo EPI on a Siemens Prisma_fit 3 T scanner (TR = 2000 ms, TE = 30 ms, FA = 90°, 37 axial slices, 3 mm isotropic voxels, 0.75 mm gap). A 1 mm isotropic T1-weighted MPRAGE and a dual-echo field map were also acquired. fMRIPrep 25.0.0 (4) ran its default workflows: FreeSurfer surface reconstruction (5), boundary-based co-registration and single-step resampling to fsLR 91k grayordinates (6). Motion parameters, framewise displacement and DVARS came from its confound output; frames above 0.5 mm framewise displacement or 1.5 standardized DVARS were flagged as outliers. Task runs entered all GLM and state-space analyses. The two resting-state runs served only as the reference frame for the gradient analysis. Regions of interest came from the combined HCP-MMP and Cole-Anticevic parcellation (7, 8). The gradient analysis used all 718 CAB-NP parcels. The ROI, state-space and memory analyses used 23 a priori target regions (162 parcel units): V1–V4; LO1–LO3 and PH; PHA1–PHA3; anterior and posterior hippocampus (split at MNI $y = -21$), amygdala and caudate; default-network PCC, mPFC and lateral temporal cortex; frontoparietal dlPFC and PPC; dorsal-attention FEF, IPS and SPL.

##### Time-lagged GLM (Fig. 4c)

A finite impulse response (FIR) GLM centered on the primary surprise event covered lags $-4$ to $+9$ TR with stick regressors and no canonical HRF. The $+2$-, $+3$- and $+4$-TR windows named throughout are BOLD sampling windows, not neural stages. The continuous-surprise model had six terms: a non-surprise viewing reference, the FIR main effect of boundary events, the FIR parametric term for the surprise score, events of no interest, six motion parameters and a constant. The high-low model split the reference term and the main effect by high- and low-surprise videos and had no parametric term. First-level data were smoothed at 4 mm FWHM. Group inference used PALM (9) with 5,000 sign-flipping permutations, threshold-free cluster enhancement (10) and FWE correction at $p < 0.05$ (minimum extent 10 $mm^2$ or 10 voxels). Table S1 gives the peak statistics and Figs. S1 and S2 the maps for all windows. Table S2 gives the ROI time courses, with grayordinates averaged within each ROI and one-sample and paired $t$ tests across participants at each lag.

##### Video-wise LSS-FIR GLM (Fig. 4a,b,d)

A least-squares-separate FIR GLM (LSS-FIR; 11) was fitted per video and participant (1,764 models). Each model coded the target video and the pooled other 35 videos over lags $-4$ to $+9$ TR, and only the target coefficients were kept. Nuisance regressors were run-specific: 24 motion columns, a constant, linear and quadratic drift, two HRF-convolved non-primary event columns, and one column per volume with framewise displacement above 0.5 mm. Models were fitted on unsmoothed grayordinate data by ordinary least squares after residualization against the nuisance set, and coefficients were converted to percent signal change. The 417 non-estimable target coefficients (1.69%, in rank-deficient models, almost all coinciding with a motion-spike column) were stored as missing. Coefficients were standardized with the trial-level participant parameters and projected into the frozen trial-level PCA of the 23-ROI space, so Figs. 3 and 4 share one coordinate system and one non-surprise reference. Because basis and nuisance treatment differ from the trial-wise GLM, the two sets of raw coefficients are not on a common scale. Group trajectories are condition-by-lag means; the lines in Fig. 4a are interpolations for display. The high-surprise trajectory was farthest from the reference at lag $+3$ and the low-surprise trajectory at lag $+2$. The high-surprise route was the more circuitous (path length over net displacement 2.64 versus 2.30). These two properties are descriptive. Temporal clusters of the high-minus-low displacement $\Delta$PC1 were formed from the paired $t$ values, joining contiguous same-sign lags, with the summed $t$ as statistic and 20,000 sign-flip permutations. High exceeded low at lags $-1$ to 0 ($p = 0.044$), $+3$ to $+4$ ($p = 0.035$; $d_z = 0.47$, $t(48) = 3.28$) and $+9$ ($p = 0.039$; Fig. S4). The lag $+9$ cluster falls on the last modeled lag and is not

**Table S1.** Peak statistics of the time-lagged GLM by peri-event window. Values are $t(48)$ at the cluster peak, TFCE-FWE $p < 0.05$. Main effects are from the condition-collapsed model, in which high- and low-surprise trials enter a single boundary-event term; condition maps and the parametric term are from the two models described above. L/R, hemisphere. Main-text Fig. 4c shows the high-surprise maps at $0$, $+3$ and $+6$ TR next to the non-surprise viewing term; Figs. S1 and S2 show all windows and models.

| Window | Term | Peaks, $t(48)$ |
|---|---|---|
| Reference | main effect | R MT 19.71, L MT 18.56; L V1 −17.94, R V1 −17.88; posterior medial and retrosplenial cortex negative in both conditions |
| +2 TR | main effect | L FST 14.31, R AVI 10.82, L PFt 10.80; R 7m −11.63, L v23ab −11.48 |
| +3 TR | main effect | L FST 14.32, L MI 12.04, R AVI 10.77 |
| +4 TR | main effect | L V1 13.20, R V1 11.88, L MI 11.66, R caudate 10.40 |
| −4 to +9 TR | high − low | after the annotated moment only visual clusters survive, all low > high: R VMV3 −8.01, R V2 −8.85, L V1 −8.97; before it, one L 7Am cluster (high > low) |
| +2 / +3 / +4 TR | high-surprise map | R AVI 9.63 / L AIP 9.10 / L 7Pm 10.55, R IP2 10.50 |
| +2 / +3 / +4 TR | low-surprise map | R VMV3 9.68 / L V4 8.72, R PH 8.17 / L V1 11.40, R V2 9.91 |
| +2 / +3 / +4 TR | parametric (rating) | R VVC 6.69 / R VMV3 8.90, R IPS1 7.69, R IP0 6.57, R 6r 6.07, R IFJp 5.94, R PGp 5.73 / R V2 6.68 |

**Table S2.** ROI time courses from the time-lagged GLM: $t(48)$ at the $+2$-, $+3$- and $+4$-TR windows. One-sample tests against zero for the condition responses and the parametric term; paired tests for high − low (negative values, low > high). Dashes, not significant; where only the significance level was recorded it is given in place of $t$.

| Contrast | ROI | +2 TR | +3 TR | +4 TR |
|---|---|---|---|---|
| Low-surprise response | V4 | 4.65 | 7.99 | 6.99 |
| | PHA3 | 4.39 | 5.51 | 3.47 |
| | V1 | — | 5.50 | 8.52 |
| | V2 | — | 3.90 | 7.40 |
| High − low | V4 | −3.82 | −4.77 | −3.98 |
| | PHA3 | −3.13 | −3.36 | — |
| | V1, V2 | — | $p < 0.010$ | $p < 0.010$ |
| High-surprise response | angular gyrus | 2.73 | 8.15 | 10.21 |
| | dlPFC | — | — | 3.00 |
| Parametric (rating) | V4 | 4.45 | 6.16 | 4.80 |
| | PHA3 | 5.08 | 4.99 | 3.01 |
| | V1, V2 | — | $p \leq 0.006$ | $p \leq 0.006$ |

interpreted. Network and parcel contributions to PC1 (Fig. 4d, Figs. S3 and S5) come from the whole-brain decomposition and support orderings, not absolute values.

#### Trial-wise GLM and state space (Figs. 2, 3, 5)

The two-step GLMsingle-style procedure of the main text (12) first selected the HRF from 20 candidates per grayordinate and the noise components explaining 50% of noise variance. It then fitted a 37-column single-video design (18 high-surprise videos, 18 low-surprise videos, events of no interest) with within-run polynomial trends and those components. Surprise designs used the primary surprise event; non-surprise designs used the video onset. Each video thus contributes one non-surprise and one surprise estimate. The two are estimated in separate designs without mutual adjustment, and the non-surprise estimate is anchored at video onset, so position within the video is confounded with state (main text, Limitations). Betas were converted to percent signal change and clipped to $[-100, 100]$. The trial-level matrix had rows for participant $\times$ condition $\times$ video and columns for the 23 ROI means, $z$-scored across the 72 observations within each participant and feature. Participant-level states are within-participant means of the low-dimensional scores. PCA was fitted jointly on the states compared, with two components for the state space and three for trajectories. PC1 was oriented to correlate positively with the surprise score at the video level and to make the mean surprise-minus-reference displacement positive at the participant level. UMAP (13) (20 neighbors, minimum distance 0.001, cosine metric, precompression to 50 components) was fitted jointly on both states over all grayordinates and used for visualization only (Fig. 2a).

#### Macroscale functional gradients (Fig. 2b–g)

Gradients were estimated with BrainSpace (14, 15) from the grayordinate-by-grayordinate Pearson correlation across single-video betas within each participant and task state; resting-state connectivity came from the denoised time series. The embedding used a normalized-angle kernel, sparsity 0.9 and 10 components, with Procrustes alignment to the pre-task resting state. Along Gradient 1 the visual and default-mode centroids lie at $-0.083$ and $+0.108$ (unimodal negative, transmodal positive). Along Gradient 2 the visual and somatomotor centroids lie at $-0.131$ and $+0.091$. For each participant and state we computed the standard deviation along each gradient, the convex-hull area, the bivariate dispersion, the eccentricity and anisotropy ratio of the covariance ellipse, and the correlation between the two gradients. States were compared by paired $t$ tests ($n = 49$). Besides the change in correlation reported in the main text ($d_z = 1.23$), the anisotropy ratio rose from 2.72 to 4.77 ($d_z = 0.65$) and the eccentricity from 0.731 to 0.834 ($d_z = 0.65$). The axis standard deviations ($d_z = 0.59$ and 0.51) and the hull area ($d_z = 0.51$) grew less. Along the a priori directions $(1,1)/\sqrt{2}$ and $(-1,1)/\sqrt{2}$ (variance $u^\top \Sigma u$), the distribution stretched along the diagonal (median $+38.8\%$, $t(48) = 7.88$, $p = 3.3 \times 10^{-10}$, $d_z = 1.13$) and compressed across it (median $-16.9\%$, $t(48) = -4.34$, $p = 7.2 \times 10^{-5}$, $d_z = -0.62$). The centroid moved by less than 0.001. These are the values annotated on Fig. 2d; network-level displacements are in Table S3.

**Table S3.** Participant-level network displacement along the first two gradients (non-surprise to surprise, $n = 49$). Positive values on Gradient 1 are toward the transmodal end and on Gradient 2 toward the somatomotor end. $q$ is FDR-corrected across the 24 tests in this table (12 networks $\times$ 2 gradients).

| Network | $\Delta G$ | 95% CI | $d_z$ | $q$ |
|---|---|---|---|---|
| *Gradient 1: unimodal (−) to transmodal (+)* | | | | |
| Visual1 | $+0.0089$ | $[+0.0066, +0.0111]$ | $+1.13$ | $2.2 \times 10^{-9}$ |
| Cingulo–Opercular | $+0.0064$ | $[+0.0044, +0.0084]$ | $+0.93$ | $2.6 \times 10^{-7}$ |
| Auditory | $+0.0056$ | $[+0.0030, +0.0082]$ | $+0.62$ | $1.6 \times 10^{-4}$ |
| Orbito–Affective | $+0.0042$ | $[+0.0026, +0.0058]$ | $+0.76$ | $6.4 \times 10^{-6}$ |
| Frontoparietal | $+0.0038$ | $[+0.0024, +0.0052]$ | $+0.77$ | $6.0 \times 10^{-6}$ |
| Somatomotor | $+0.0033$ | $[+0.0007, +0.0059]$ | $+0.37$ | 0.017 |
| Ventral–Multimodal | $+0.0011$ | $[-0.0007, +0.0029]$ | $+0.18$ | 0.258 |
| Default | $+0.0003$ | $[-0.0016, +0.0022]$ | $+0.05$ | 0.717 |
| Language | $-0.0018$ | $[-0.0039, +0.0003]$ | $-0.24$ | 0.125 |
| Posterior–Multimodal | $-0.0035$ | $[-0.0052, -0.0018]$ | $-0.61$ | $2.0 \times 10^{-4}$ |
| Visual2 | $-0.0044$ | $[-0.0067, -0.0021]$ | $-0.54$ | $6.7 \times 10^{-4}$ |
| Dorsal–Attention | $-0.0051$ | $[-0.0073, -0.0028]$ | $-0.65$ | $9.2 \times 10^{-5}$ |
| *Gradient 2: visual (−) to somatomotor (+)* | | | | |
| Visual1 | $+0.0109$ | $[+0.0089, +0.0129]$ | $+1.54$ | $4.6 \times 10^{-13}$ |
| Default | $+0.0069$ | $[+0.0052, +0.0086]$ | $+1.17$ | $1.3 \times 10^{-9}$ |
| Orbito–Affective | $+0.0050$ | $[+0.0031, +0.0068]$ | $+0.78$ | $5.6 \times 10^{-6}$ |
| Frontoparietal | $+0.0042$ | $[+0.0027, +0.0057]$ | $+0.80$ | $4.6 \times 10^{-6}$ |
| Somatomotor | $+0.0041$ | $[+0.0019, +0.0063]$ | $+0.53$ | $8.2 \times 10^{-4}$ |
| Auditory | $+0.0039$ | $[+0.0010, +0.0069]$ | $+0.38$ | 0.015 |
| Ventral–Multimodal | $+0.0032$ | $[+0.0013, +0.0052]$ | $+0.48$ | 0.002 |
| Cingulo–Opercular | $+0.0008$ | $[-0.0009, +0.0026]$ | $+0.14$ | 0.373 |
| Language | $+0.0003$ | $[-0.0013, +0.0019]$ | $+0.05$ | 0.717 |
| Visual2 | $-0.0011$ | $[-0.0036, +0.0013]$ | $-0.13$ | 0.396 |
| Dorsal–Attention | $-0.0038$ | $[-0.0057, -0.0019]$ | $-0.58$ | $3.5 \times 10^{-4}$ |
| Posterior–Multimodal | $-0.0040$ | $[-0.0053, -0.0027]$ | $-0.88$ | $6.9 \times 10^{-7}$ |

#### Memory models (Fig. 5)

Overall exceedance is each participant's mean over all videos; condition-specific exceedance and memory were computed separately for high- and low-surprise videos. Moderation used random-intercept mixed-effects models (REML, Satterthwaite degrees of freedom), with ordinary least squares and participant-clustered robust standard errors as the fallback for singular fits. The moderation was not predicted and is reported as exploratory; the primary interaction survives Bonferroni correction across the three recognition outcomes (accuracy, reaction time, log reaction time; $p = 0.00085$, corrected $p = 0.0026$). The low-minus-high model regressed each participant's low- minus high-surprise performance on overall exceedance.

#### Boundary exceedance: definition, invariance and sensitivity

The construction is in the main text. Below, $\hat{\mathbf{u}}$ is the transition axis and $R$ the reference population of all non-surprise trials. For $X \sim R$, $T = \hat{\mathbf{u}}^\top X$ is the reference projection and $\pi(x) = \Pr(T \geq \hat{\mathbf{u}}^\top x)$ the upper-tail probability of a state $x$. The axis, the reference population and the contour probability $q$ must all be stated for a value in bits to be interpretable. Because $D = -\log_2 \pi$

is non-linear in $\pi$, the depth of a mean state is not the mean of depths; the video-level points in Fig. 3c are means of trial-level depth.

*Invariance.* $\pi$ depends on the projected coordinate only through its rank. $E_{\mathrm{BB}}$ is therefore invariant to any strictly increasing transformation along the axis and to isotropic rescaling of the state space; multiplying all coordinates by 10 changed nothing, while $\|\Delta\mathbf{S}\|$ rose from 1.985 to 19.845. It is not invariant to anisotropic rescaling, because the mean-difference axis is defined in a particular metric. Scaling PC2 by 0.5 or 1.5 rotated the axis by about 7° and changed mean breakthrough exceedance by about 11% (3.415 bits to 3.018 and 3.795). Scaling by 3 or 6 rotated it by 23° and 43° and cut exceedance by 37% and 80%.

*Contour probability.* For states beyond the stricter of two boundaries, $E_{\mathrm{BB}}(x;q) - E_{\mathrm{BB}}(x;q') = \log_2(q/q')$. Changing $q$ therefore shifts the scale rigidly and alters only which states are censored. Over $q \in [0.01, 0.20]$ the video-level correlation with rated surprise stays between $r = 0.402$ and 0.421, the paired slope difference on $E_{\mathrm{BB}}$ stays significant ($p \le 0.016$), and the slope difference on $D$ is $\beta = +0.564$ bits per $z$ at every $q$.

*Transition axis.* The axis is estimated from the contrast it serves, so every quantity was recomputed under four alternatives (Table S4). All give a positive correlation and slope difference, and the axis is estimated precisely (leave-one-video-out rotation $\le 0.87°$). The Fisher-discriminant and whitened axes do not reach significance. The effect is carried by displacement along PC1, which both down-weight; the first principal axis alone gives the strongest result, so the mean-difference axis does not maximize the surprise association. The axes that best separate the conditions track rated surprise least, so the beyond-boundary rate is no measure of validity (the reference rate is 4.99% under every axis by construction). We keep the mean-difference axis because the framework specifies it and no outcome was used to choose it. Its composition is in Table S5.

**Table S4.** Sensitivity of the reported quantities to the choice of transition axis. Angles are relative to the mean-difference axis. The video-level correlation is between mean $E_{\mathrm{BB}}$ and rated surprise ($n = 36$); the paired slope difference is the primary test reported in main-text Fig. 3c. Ceiling is the proportion of breakthrough trials at the empirical maximum.

| Axis | angle | breakthrough beyond | $r(E_{\mathrm{BB}}, \text{rating})$ | paired $\Delta$slope | ceiling |
|---|---|---|---|---|---|
| mean difference (used here) | — | 79.6% | +0.419 ($p = 0.011$) | +0.564 ($p = 0.015$) | 23.0% |
| Fisher discriminant | 30.1° | 87.2% | +0.247 ($p = 0.146$) | +0.279 ($p = 0.196$) | 27.6% |
| whitened mean difference | 21.3° | 86.3% | +0.315 ($p = 0.062$) | +0.369 ($p = 0.085$) | 25.3% |
| first principal axis | 14.6° | 74.9% | +0.442 ($p = 0.007$) | +0.646 ($p = 0.009$) | 22.4% |
| leave-one-video-out | 0.25° | 79.6% | +0.421 ($p = 0.011$) | +0.569 ($p = 0.015$) | 23.2% |

*Finite-sample estimation and the ceiling.* $\hat{\pi}(x) = \max\{k(x)/n,\ 1/(n+1)\}$, with $k(x)$ the number of reference trials at or beyond $x$, floors the estimate. The add-one estimator moves mean breakthrough exceedance from 3.415 to 3.250 bits and changes no conclusion ($r = 0.418$ versus 0.419; paired slope difference $\beta = +0.431$ versus $+0.440$, both $p = 0.012$). The scale is discrete, with steps widening from 0.002 bits near the reference median to a full bit at the 6.464-bit ceiling, so the deepest bit is ordinal. Four checks show that the surprise scaling does not depend on the 23.0% of breakthrough trials at the ceiling. Rank statistics agree with the Pearson correlation ($r(34) = 0.419$, $p = 0.011$; Spearman $\rho = 0.433$, $p = 0.008$; Kendall $\tau = 0.305$, $p = 0.009$). A trial-level Tobit regression censored at 6.464 bits gives a larger coefficient than ordinary least squares ($\beta = +0.575$ versus $+0.446$ bits per $z$), the direction expected if censoring attenuates a monotonic relation. A generalized Pareto fit above the 90th percentile of the reference projection removes the ceiling and keeps the association ($r(34) = 0.382$, $p = 0.021$). The 18 videos with below-median ceiling proportion give the same point estimate ($r(16) = 0.423$, $p = 0.081$).

##### Construction checks

*The transform.* $D = -\log_2 \pi$ expands the thin end of the reference distribution. The local gain $\mathrm{d}D/\mathrm{d}v = f(v)/[\pi(v)\ln 2]$ is 1.64 and 5.69 bits per PC unit at the two condition centroids ($v = \mp 0.99$), so part of the difference between the rating slopes could come from the transform. The main text repeats the contrast on the untransformed axis position, where both states share one linear scale, with the same inference. *Cross-fitting.* For every video, the axis and the trial-level reference distribution were re-estimated from the other 35 videos and applied to the held-out video; the label permutation repeated that pipeline (Table S4). The PCA basis was fitted once on all trials, so the check covers axis and boundary, not the state space. *Pooling across participants.* Centering every trial on its own participant's reference mean before re-estimating the boundary left the beyond-boundary rate at 80.4% (79.6% pooled) and the exceedance–rating association at $r = 0.43$, $p = 0.010$. For the geometry–memory relation the boundary was estimated from the other 48 participants and applied to each held-out participant.

#### Supplementary Results

##### State-specificity of the surprise scaling

The rating × state model of the main text was fitted on boundary depth $D$ with a random intercept per video ($n = 36$ paired videos). It uses $D$ rather than $E_{\mathrm{BB}}$ because exceedance is censored at zero for 95% of reference trials by construction. At the video level the reference-state $E_{\mathrm{BB}}$ is $0.070 \pm 0.087$ bits against $1.439 \pm 0.293$ bits for $D$, so a comparison on $E_{\mathrm{BB}}$

**Table S5.** Composition of the transition axis. Loadings on the first two principal components and the resulting axis weight $w$, ordered by $w$. Shares of $\sum|w|$: dorsal-attention and frontoparietal $41.2\%$ (signed sum $+1.577$), LO1–LO3 and PH $30.1\%$ ($+1.150$), medial temporal $9.4\%$ ($-0.327$), default-mode $9.3\%$ ($-0.059$), early visual V1–V4 $7.5\%$ ($+0.229$), caudate $2.5\%$ ($+0.095$). Fifteen of the weights are positive and the cosine between the axis and the uniform vector is $+0.556$, so the axis carries a general positive component alongside the control-versus-medial-temporal contrast.

| Region | PC1 | PC2 | $w$ |
|---|---|---|---|
| Dorsal-attention IPS | $+0.367$ | $-0.293$ | $+0.429$ |
| Dorsal-attention SPL | $+0.353$ | $-0.315$ | $+0.421$ |
| PH | $+0.361$ | $-0.029$ | $+0.357$ |
| Dorsal-attention FEF | $+0.267$ | $-0.238$ | $+0.318$ |
| LO1 | $+0.382$ | $+0.379$ | $+0.274$ |
| Frontoparietal PPC | $+0.213$ | $-0.250$ | $+0.270$ |
| LO3 | $+0.343$ | $+0.269$ | $+0.264$ |
| LO2 | $+0.333$ | $+0.262$ | $+0.256$ |
| Default lateral temporal | $+0.143$ | $-0.039$ | $+0.148$ |
| Frontoparietal dlPFC | $+0.102$ | $-0.164$ | $+0.140$ |
| V4 | $+0.208$ | $+0.307$ | $+0.123$ |
| Caudate | $+0.071$ | $-0.102$ | $+0.095$ |
| V3 | $+0.126$ | $+0.176$ | $+0.077$ |
| V1 | $+0.082$ | $+0.088$ | $+0.057$ |
| Amygdala | $+0.023$ | $+0.022$ | $+0.017$ |
| V2 | $+0.011$ | $+0.154$ | $-0.028$ |
| Anterior hippocampus | $-0.020$ | $+0.064$ | $-0.036$ |
| Posterior hippocampus | $-0.023$ | $+0.066$ | $-0.039$ |
| PHA3 | $+0.007$ | $+0.222$ | $-0.050$ |
| Default mPFC | $-0.068$ | $+0.076$ | $-0.085$ |
| PHA2 | $-0.034$ | $+0.270$ | $-0.101$ |
| PHA1 | $-0.063$ | $+0.226$ | $-0.118$ |
| Default PCC | $-0.084$ | $+0.163$ | $-0.122$ |

would set a slope against a floor. The random-intercept variance was estimated at zero, so the model reduces to ordinary least squares with 68 residual degrees of freedom. Because the states differ markedly in dispersion (residual SD 0.288 versus 1.209 bits), coefficients carry HC3 robust standard errors, with $t(68)$ inference for $p$ values and confidence intervals. The 95% CIs are $[0.103, 1.010]$ for the breakthrough-state slope ($p = 0.017$), $[-0.131, 0.116]$ for the reference-state slope ($p = 0.90$) and $[0.094, 1.034]$ for the interaction ($p = 0.020$); clustering by video gives essentially the same $p$. Indexing surprise by the single normative surprise rating instead of the factor score, which correlates with it at $r = 0.99$ across videos, gave a paired slope difference of $\beta = 0.555$ bits per s.d., $t(34) = 2.47$, $p = 0.019$, and an exceedance–rating correlation of $r(34) = 0.41$, $p = 0.014$. The paired video-level test of the main text has 95% CI $[0.116, 1.012]$. On $E_{\mathrm{BB}}$ rather than $D$ it gives $\beta = 0.440$, $t(34) = 2.67$, $p = 0.012$. For the direct exceedance–rating correlation, a bootstrap over videos (10,000 resamples) gave a 95% CI of $[0.10, 0.66]$ and a label permutation (10,000 shuffles) gave $p = 0.010$. The Euclidean magnitude correlates more weakly and not significantly ($r(34) = 0.306$, $p = 0.070$), as expected when along-axis and orthogonal movement are mixed.

#### Transition direction and curiosity controls

The direction-given-depth effect reported in the main text for the low-over-high recognition advantage also held for low-surprise recognition accuracy. Adding $\theta = \mathrm{atan2}(\Delta\mathrm{PC2}, \Delta\mathrm{PC1})$ to a model containing $E_{\mathrm{BB}}$ gave $\Delta R^2 = 0.136$, $F(1, 46) = 8.22$, $p = 0.006$. With the Euclidean magnitude in place of $E_{\mathrm{BB}}$ the gain is smaller but concordant ($\Delta R^2 = 0.085$, $p = 0.023$). The angle alone was unrelated to recognition ($r = -0.09$, $p = 0.56$). The contribution survived trait curiosity, state curiosity and need-for-cognition as covariates (all $p \leq 0.010$), and $E_{\mathrm{BB}}$ was uncorrelated with them (all $p > 0.37$). Reward value was not modeled and covaries with surprise in these materials. These analyses span four recognition outcomes without correction for multiple comparisons and are exploratory.

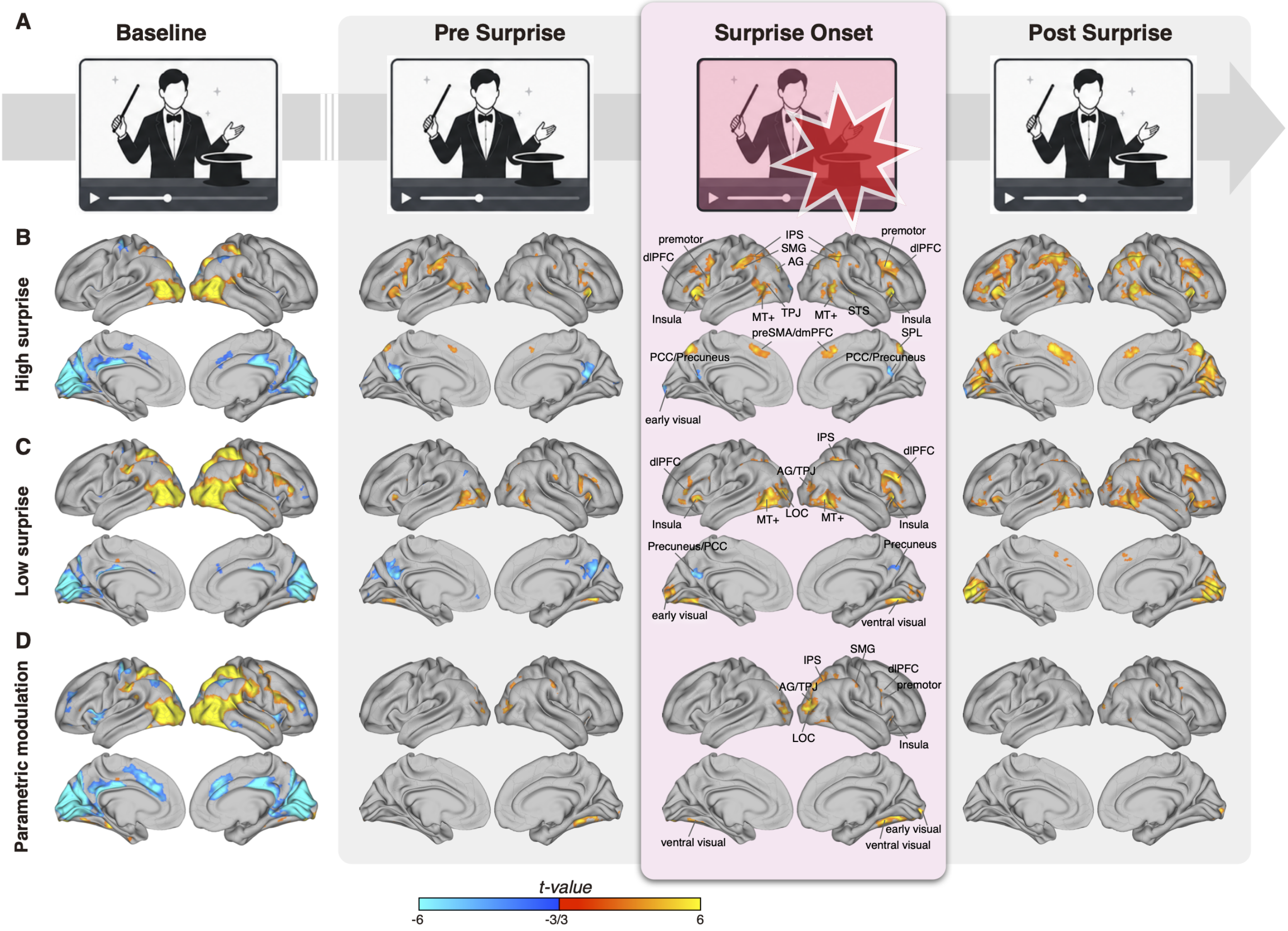


**Fig. S1.** Cortical time-lagged GLM results across the peri-event window, both hemispheres (main-text Fig. 4c shows the left hemisphere of row **B** at three windows, with the baseline). Columns are BOLD sampling windows relative to the surprise marker. **A**, Temporal structure of a trial. **B**, High-surprise condition. **C**, Low-surprise condition. **D**, Continuous surprise-intensity modulation. Peak regions are labeled in the shaded surprise-onset column; colors give $t(48)$, warm positive and cool negative.

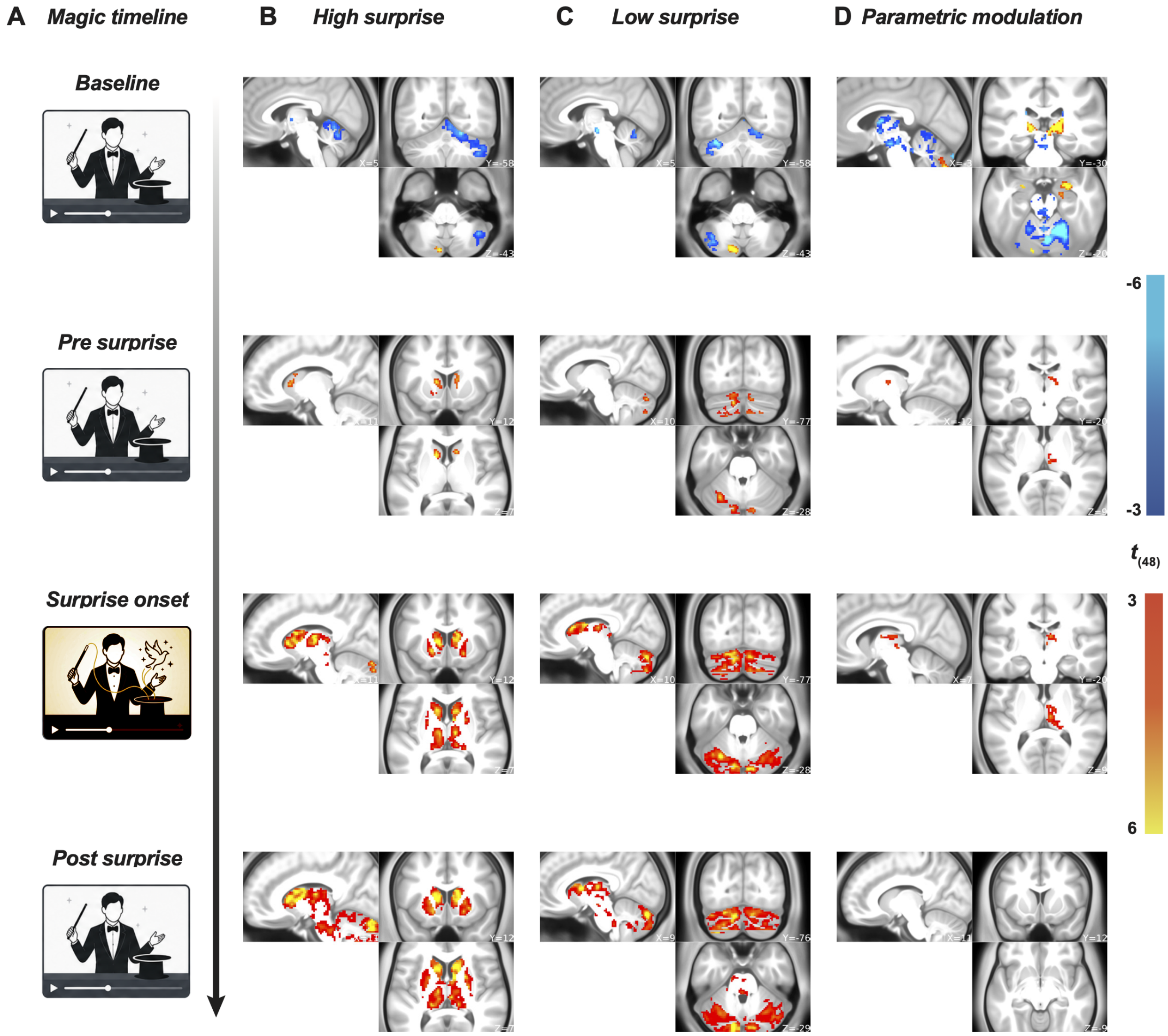


**Fig. S2.** Subcortical time-lagged GLM results for the $+2$-, $+3$- and $+4$-TR windows after the annotated moment. **A**, Temporal structure of a trial. **B**, High-surprise condition. **C**, Low-surprise condition. **D**, Continuous surprise-intensity modulation. Colors give $t(48)$, warm positive and cool negative.

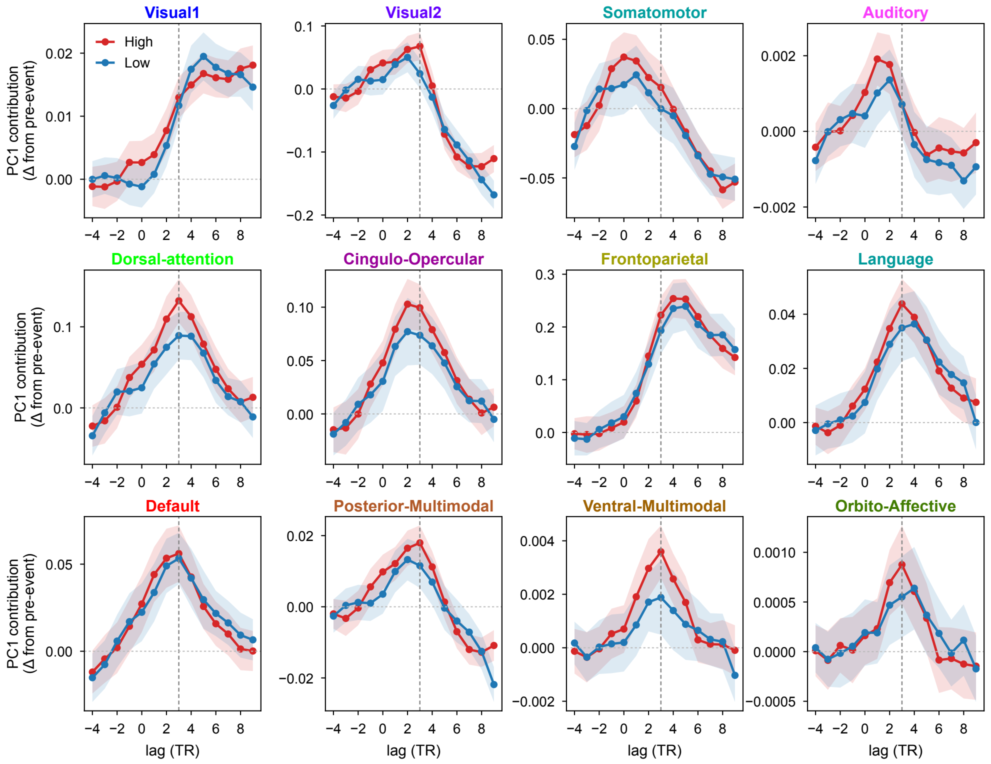


**Fig. S3.** Contribution of every Cole-Anticevic network to PC1 across the peri-event window (whole-brain PCA; red, high surprise; blue, low surprise; bands, bootstrap 95% CIs over participants, $n = 49$; curves referenced to the pre-event baseline, lags $-4$ to $-1$; dashed line, the $+3$-TR window). Main-text Fig. 4d shows the two networks with the most extreme post-event value; this is the full set they were taken from. The networks differ mainly in where they end up rather than when they peak: only Frontoparietal remains clearly above baseline late in the window, whereas Visual2 and Somatomotor fall below it. Vertical scales differ across panels.

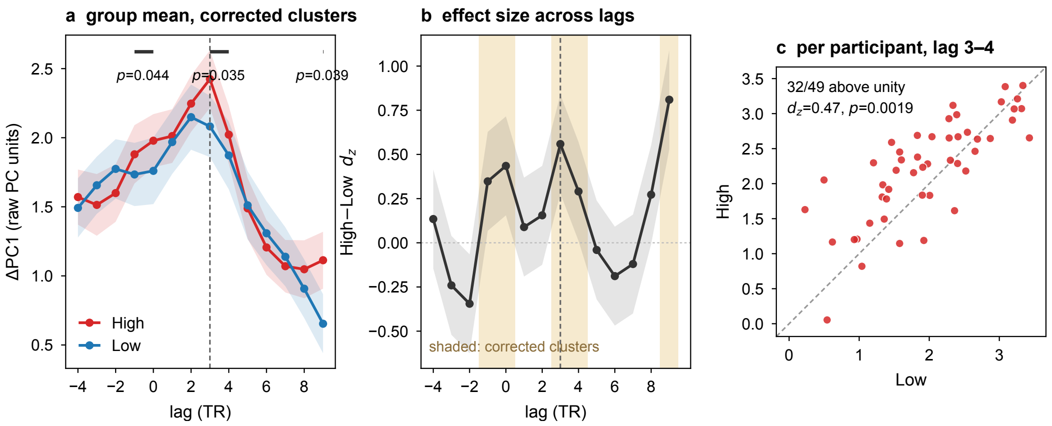


**Fig. S4.** Lag-resolved directional displacement ΔPC1, in raw PC units of the frozen trial-level PCA of Figs. 3 and 4, referenced to the same non-surprise state. **a**, Group mean per lag for high- and low-surprise events (shading, ±1.96 SEM; $n = 49$); bars mark the FWER-corrected high-versus-low clusters with their $p$ values. **b**, High−low effect size $d_z$ across lags; the effect peaks at the $+3$-TR window. **c**, Participant values over lags $+3$ to $+4$ against the identity line. The lag $+9$ cluster falls on the last modeled lag and is not interpreted.

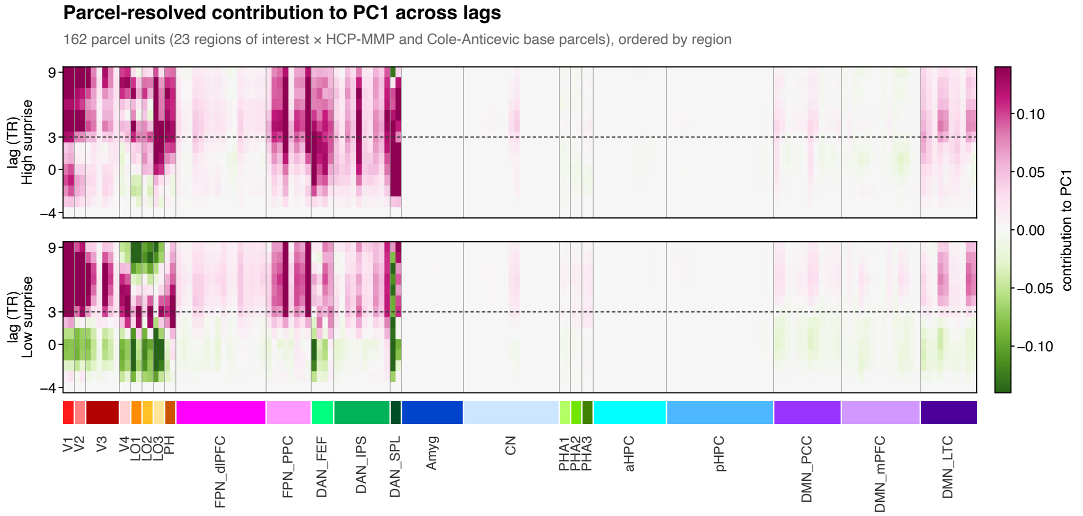


**Fig. S5.** Parcel-resolved contribution to PC1 across lags for the 162 parcel units, ordered by region (color strip below). Upper band, high-surprise events; lower band, low-surprise events; dashed line, the $+3$-TR window. At this scale the conditions separate by sign rather than magnitude: lateral-occipital and superior parietal parcels contribute positively under high surprise but negatively under low surprise (LO1, $+0.070$ versus $-0.095$; SPL, $+0.107$ versus $-0.013$), frontoparietal parcels positively in both, and subcortical parcels negligibly. Contributions are to PC1 of the whole-brain decomposition and support the ordering of parcels rather than absolute values.
